# Genomic signatures and demographic history of the widespread and critically endangered yellow-breasted bunting

**DOI:** 10.1101/2024.08.26.609693

**Authors:** Guoling Chen, Simon Yung Wa Sin

## Abstract

Population declines may have long-term genetic consequences, including genetic erosion and inbreeding depression, which could affect species’ evolutionary potential and increase their risk of extinction. Small populations are more vulnerable to genetic threats than common species, but even species with large populations can also be at risk of extinction. The yellow-breasted bunting (*Emberiza aureola*) is a common and widespread songbird in the northern Palearctic regions, but its population has drastically declined by around 90% throughout the past 30 years, leading to an upgrade of its conservation status to critically endangered. In this study, we identified three populations within this species using whole-genome resequencing data, but the genetic differentiation between populations was shallow. These populations underwent similar population fluctuations but differed in the extent of population decline, resulting in lower genetic diversity and more homozygous deleterious mutations in a population comprising individuals on islands. The ancient demographic history was mainly associated with the climate, while recent population declines are likely due to human activities. Our results suggest that the yellow-breasted bunting population before the recent population collapse faced relatively low genetic threats and had high evolutionary potential. However, we should be vigilant about the genetic threats faced by this species as our sampling time occurred at the onset of its recent global population collapse. This study provides valuable genetic information for appropriate conservation management of the yellow-breasted bunting and sheds light on the extinction risks and genetic consequences common species face.

## 1 Introduction

Many species are undergoing continuous population decline attributable to both climatic change and human activities (Cowie et al., 2022). Fluctuations in ancient climate have significantly impacted the population of species (Nadachowska-Brzyska et al., 2015). Many species underwent distribution range contraction and fragmentation during glacial periods, while interglacial periods facilitated population expansion (Hewitt, 2004; Holm & Svenning, 2014; Nadachowska-Brzyska et al., 2015). In recent history, human activities have had an increasingly significant impact on biodiversity, emerging as one of the main causes of population decline and species extinction. Currently, more attention is being given to ecological and anthropogenic factors to prevent biodiversity loss and population decline. However, genetic factors are often neglected due to the challenges in accessing and applying them directly to conservation management, despite their importance in species conservation (Kardos et al., 2016).

Population decline can result in various genetic consequences, including the loss of genetic diversity (Frankham et al., 2010; Gandra et al., 2021; Huynh et al., 2023). Genetic diversity is pivotal in a species’ ability to adapt to changing environments. The loss of genetic diversity in a species can have a significant impact on its long-term evolutionary potential (Kardos et al., 2021). Another possible consequence of population decline is inbreeding and subsequent inbreeding depression (Sin et al., 2021). Population decline may increase inbreeding, leading to inbreeding depression due to the exposure of harmful mutations in homozygous state (Hedrick & Garcia-Dorado, 2016). Previous studies suggest that strong deleterious mutations are the main determinants of inbreeding depression, which may increase the risk of extinction (Kyriazis et al., 2021). Additionally, the number of these mutations in a population often correlates with the effective population size (Bertorelle et al., 2022). Smaller populations that persist for a long time tend to have more homozygous deleterious mutations, possibly due to the increase in inbreeding and genetic drift and less effective purifying selection in a small population (Bertorelle et al., 2022). In contrast, larger populations generally harbor more heterozygous deleterious mutations (Kyriazis et al., 2021). Consequently, larger populations are more susceptible to inbreeding depression during population declines. During a population decline, some deleterious mutations may be eliminated by purifying selection, while others may become fixed and negatively impact the overall fitness of the population (Glemin, 2003; Grossen et al., 2020). Therefore, assessing genetic features like genetic diversity, inbreeding level, and genetic load can provide valuable insights into the current genetic health and future viability of threatened species.

Endangered species, particularly those with small populations, often exhibit low genetic diversity and high inbreeding, which renders them more vulnerable to the “extinction vortex” (Blomqvist et al., 2010; Fagan & Holmes, 2006). As a result, small populations usually receive more attention in conservation efforts than common species with large populations. Currently, the majority of research in conservation genetics focuses on small populations, such as the vaquita, killer whales, and Isle Royale wolves (Kardos et al., 2023; Robinson et al., 2022; Robinson et al., 2019). In contrast, we may overlook population declines in common species despite their important role in maintaining ecosystem balance (Inger et al., 2015). Previous studies indicate that common European birds, such as the house sparrow and common starling, experienced more significant population declines compared to less abundant species (De Laet & Summers-Smith, 2007; Inger et al., 2015; Smith et al., 2012).

Indeed, large populations can also be at risk of extinction when faced with dramatic population fluctuations, like the passenger pigeon. In the early and mid-1880s, the passenger pigeon population was estimated to be 3-5 billion (Schorger, 1995). Shockingly, it took only five decades for this abundant bird to become extinct. Previous studies suggest that a combination of dramatic population fluctuations and human activities may have ultimately led to the extinction of this species (Hung et al., 2014). Additionally, another study emphasized that although larger populations like the passenger pigeon exhibited the ability to eliminate harmful mutations and had developed traits to adapt to its environment, it could still face the risk of extinction following a sudden environmental change (Murray et al., 2017).

The yellow-breasted bunting (*Emberiza aureola*), which was historically one of the most abundant songbirds of the northern Palearctic region, is now facing a rapid population collapse similar to that of passenger pigeons. They breed from northern and central Europe to far eastern Russia and Japan; most populations migrate toward the east, cross Siberia and northeast China, stop in the Yangtze Valley in China, and winter in Southeast Asia (BirdLife International, 2024). The population size of this species was estimated to be hundreds of millions in the 1980s, but its numbers have drastically declined by 84.3%-94.7% between 1980 and 2013 (Kamp et al., 2015). Additionally, some populations (e.g., in Finland) have disappeared from certain breeding areas since the 2000s (Copete & Sharpe, 2020; Tamada et al., 2017; Tamada et al., 2014). Consequently, its conservation status was upgraded from Least Concern in 2000 to Critically Endangered in 2017. This alarming situation highlights the need for conservation management to protect this species. Overhunting during the migration, agricultural intensification, and habitat destruction are believed to be the primary causes of the decline in the yellow-breasted bunting population (BirdLife International, 2024; Kamp et al., 2015). However, it remains uncertain whether other genetic factors contribute to this decline. A previous study provided some genetic information on this species (Wang et al., 2022). However, due to the limitations of the small sample size (n=10) and the collection of samples during migration, information regarding the population structure, conservation units, changes in distribution range, and demographic history of different populations remains lacking.

In this study, we used whole genome resequencing data to investigate the demographic history and genetic consequences of population declines in yellow-breasted buntings. To delineate conservation units, we first determined the population structure of this species and examined gene flow between populations. We then reconstructed the ancient and recent demographic history of different populations and investigated the potential effect of paleoclimate on suitable breeding and wintering habitats of this species. Lastly, we investigated genetic features such as genetic diversity, inbreeding, and mutation load to assesse the genetic health and evolutionary potential of the yellow-breasted bunting. Our study yields valuable insights into the mechanisms that contribute to endangerment in large populations and provides knowledge that can be used to develop effective conservation strategies for this critically endangered songbird.

## 2 Materials and Methods

### 2.1 Sample collection

We acquired tissue samples from three bunting species, which included 94 yellow-breasted bunting (*Emberiza aureola*) and two outgroups (one black-faced bunting *E. spodocephala* and one reed bunting *E. schoeniclus*) from museum collections (Table S1). The samples were collected during the breeding season from 1992 to 2004, and the sampling sites of yellow-breasted bunting covered most of their breeding areas (Figure 1b). The tissue samples were stored in absolute ethanol at −80 ℃ until DNA extraction.

**Figure 1.**
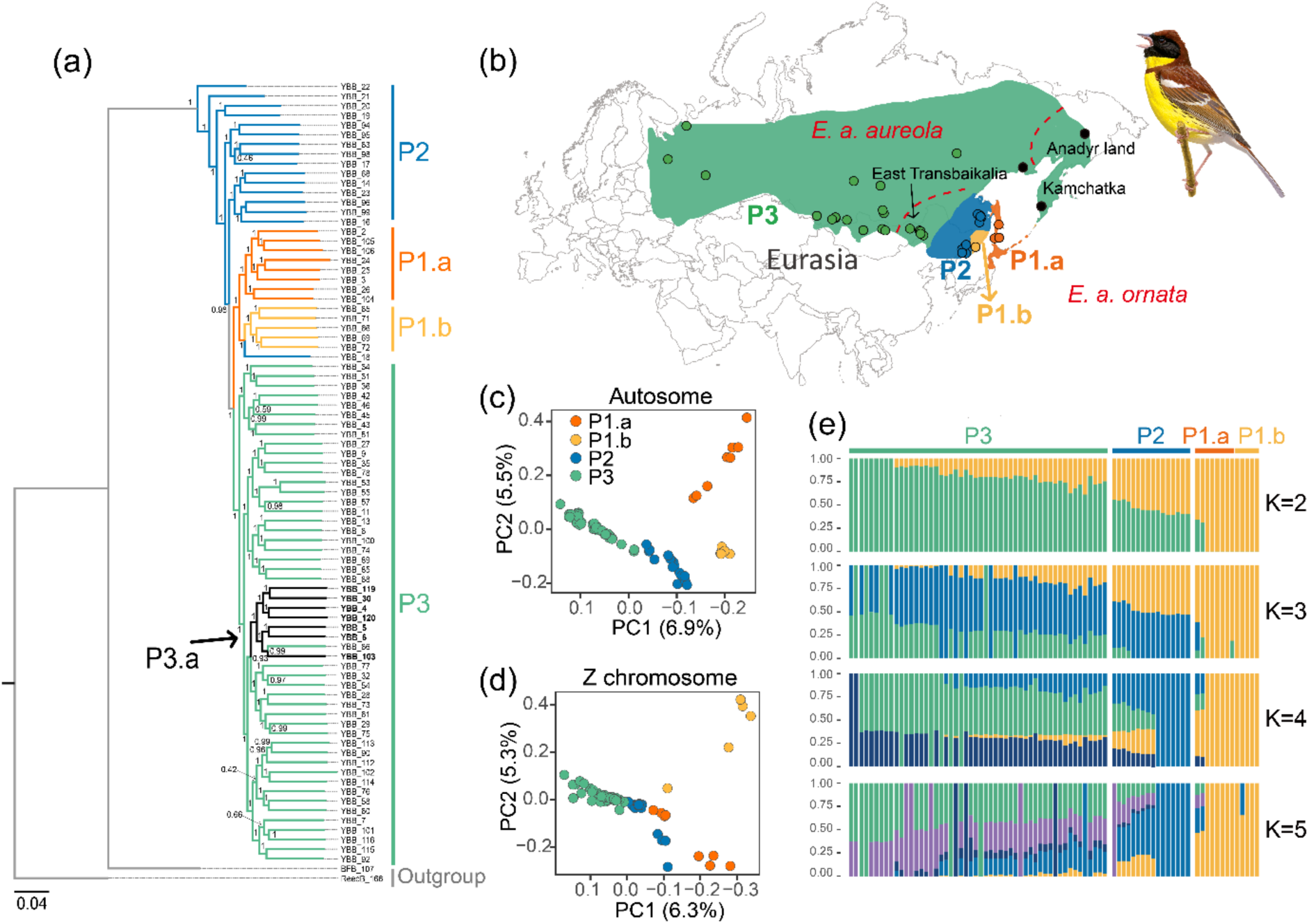
The sampling sites and population structure of yellow-breasted buntings. (a) The maximum likelihood tree was inferred using FastTree based on the autosomal SNPs. (b) The map shows the breeding area and sampling sites of yellow-breasted bunting. Different colors indicate different populations in this study. Points indicate sampling locations. The red dashed lines indicate the boundaries of the proposed subspecies distribution (Copete & Sharpe, 2020). Principal components analysis (PCA) for all the yellow-breasted bunting individuals based on (c) autosomal SNPs and (d) Z chromosomal SNPs. (e) Population structure inferred using Admixture with K ranging from 1 to 5. Each bar indicates one individual, and the y-axis indicates the probability of this individual being assigned to one or more clusters. Illustrations reproduced with permission of Lynx Edicions.

### 2.2 Genome assembly and annotation

One individual of yellow-breasted bunting was used for genome assembly (Table S1). High-quality genomic DNA was extracted from the tissue sample using the MagAttract HMW DNA Kit (Qiagen). The DNA quality was assessed using Pulse-field gel electrophoresis, and a DNA library for whole genome sequencing was prepared using Chromium (10x genomics). The library was sequenced on Illumina NovaSeq (PE 150bp reads) by Novogene (Hong Kong) to achieve 132Gb data.

We first processed the raw data by removing the low-quality data, adapters, and duplicated reads using FASTX-Toolkit v.0.0.14 (Hannon, 2010) and FastUniq (Xu et al., 2012). Then, we corrected short-read errors using Musket (Liu et al., 2013). To check the quality of both the raw and clean sequencing data, we used FastQC (Andrews, 2017). Next, we assembled the reference genome using Supernova (Zheng et al., 2016) with clean reads. To evaluate the completeness of the reference genome, we ran BUSCO (Simao et al., 2015) analysis using the Aves dataset (aves_odb10) with 4,915 universal single-copy orthologs. Finally, we mapped the scaffolds of the yellow-breasted bunting draft genome to the zebra finch (*Taeniopygia guttata*) chromosome-level genome (GCA_008822105.2) from NCBI using Satsuma (Grabherr et al., 2010) to construct the pseudochromosomes.

To annotate the reference genome, we first identified and annotated the sequences of interspersed repeat using RepeatModeler v2.0.2 (Flynn et al., 2020) and RepeatMasker v4.0.5 (Smit et al., 2020). We then train *ab initio* gene predictors with the expressed sequence tags (ESTs) using SNAP in MAKER v2.31.9 (Holt & Yandell, 2011). The ESTs were obtained from the NCBI database, comprising the protein sequences of 15 Passeriformes species. After that, the results of *ab initio* gene prediction, EST alignments, and protein alignments were combined to generate gene predictions in MAKER v2.31.9. Furthermore, homology-based gene prediction was performed based on the nine avian species in GeMoMa v1.7.1 (Keilwagen et al., 2019). Finally, the annotation results from MAKER2 and GeMoMa for the yellow-breasted bunting genome were combined using GeMoMa v1.7.1 and then evaluated using BUSCO analysis. Functional annotation for the reference genome was carried out using the UniProt/Swiss-Prot database (Bairoch & Apweiler, 2000) and eggNOG-mapper (Huerta-Cepas et al., 2019).

### 2.3 DNA extraction and whole-genome resequencing

We extracted the genomic DNA of bunting tissues using the E.Z.N.A. Tissue DNA Kit (Omega Bio-tek) following the manufacturer’s protocol. The DNA quantity and quality were assessed using the Qubit dsDNA High Sensitivity Assay Kit on the Qubit 4 fluorometer (Thermo Scientific, MA, USA) and agarose gel electrophoresis, respectively. Only high-quality DNA samples (n=94; Table S1) were used for the whole-genome resequencing. DNA libraries of 350bp insert size were prepared, and sequencing was performed on Illumina NovaSeq (PE 150bp reads) by Novogene (Hong Kong) to achieve 30Gb data per sample.

### 2.4 SNP calling and filtering

The raw data obtained from the whole-genome resequencing was filtered using Fastp (Chen et al., 2018), and the data quality was evaluated using FastQC. The clean data was then aligned to the reference genome using Burrows-Wheeler Aligner (BWA V0.5.17) (Li & Durbin, 2009) with the default settings. Subsequently, we called the single nucleotide polymorphisms (SNPs) following the GATK 4.1.4.0 (McKenna et al., 2010) Best Practices Workflows with Samtools (Li et al., 2009) and Picard (Picard, 2019).

To obtain a high-quality and credible SNP dataset, we first performed hard filtering (SOR>3, FS>60, MQRankSum < −12.5, ReadPosRankSum < −8, ReadPosRankSum > 8, MQ < 40, QD < 1) for SNPs using GATK 4.1.4.0. We then performed soft filtering using VCFtools (Danecek et al., 2011) following these criteria: --minQ 30, --minDP 5, --maxDP 70, --max-missing 0.95, --max-alleles 2. We excluded the SNPs located on the 4∼10 Mb section of chromosome 11 from all the analyses due to the higher number of missing SNPs in this region. After obtaining the results from Satsuma, we extracted the SNPs of the autosomes and the Z chromosome.

We used KING (Manichaikul et al., 2010) to calculate kinship coefficients for all pairs of individuals and excluded five individuals from our dataset with a kinship coefficient greater than 0.177 to avoid potential bias from closely related individuals. Additionally, we assessed the sequencing depth and missing rate for each individual and excluded eight individuals due to high missing rates. The remaining dataset consisted of 81 yellow-breasted buntings, one black-faced bunting, and one reed bunting that served as the outgroup for subsequent analyses.

### 2.5 Population structure and phylogenetic analysis

To avoid clustering bias caused by linkage disequilibrium (LD), we conducted LD pruning using PLINK v.1.9 (Purcell et al., 2007). For population structure analysis, we also filtered out SNPs with minor allele frequency (MAF < 0.05). We then estimated population structure using autosomal SNPs, Z chromosomal SNPs, and the mitochondrial genomes. The mitochondrial genomes were assembled using MIRA v4.0 (Chevreux et al., 2004) and MITOBIM.PL v1.6 (Hahn et al., 2013). A yellow-breasted bunting mitochondrial genome (Pan et al., 2015) available on NCBI (accession no. NC022150.1) was used as the reference for the assembly of mitochondrial genomes.

We conducted three analyses to determine the population structure of yellow-breasted buntings. First, we performed principal component analysis (PCA) using PLINK v.1.9 based on autosomal and Z chromosomal SNPs. We employed R package adegenet (Jombart, 2008; Jombart & Ahmed, 2011) to perform PCA on mitochondrial data. Second, we reconstructed the maximum likelihood (ML) tree based on autosomal SNPs using FastTree v2.1.11 (Price et al., 2010) with the General Time Reversible (GTR) model. Then, we reconstructed the ML tree using IQ-TREE2 (Minh et al., 2020) for the Z chromosomal SNPs and mitochondrial data. The best-fit substitution model for DNA sequences was determined by ModelFinder in IQ-TREE2. We used reed bunting as outgroups and performed 1000 bootstraps for each analysis with UFBoot to estimate branch support. Finally, we estimated individual admixture proportions based on autosomal SNPs using ADMIXTURE (Alexander et al., 2009). We tested genetic clusters parameter K ranging from 1 to 6 and conducted each analysis with 200 bootstraps. The best K was determined based on the value of cross-validation error.

To quantify the differentiation between populations, we estimated the relative differentiation (*F*_ST_) between the populations using VCFtools (Danecek et al., 2011). The population *F*_ST_ was calculated using 20kb non-overlapping sliding windows, and any windows with less than 50 SNPs were excluded from the analysis. Isolation by distance is a common model that illustrates the correlation between genetic divergence and dispersal among various geographic regions (Lam et al., 2023). To investigate whether the distribution of the yellow-breasted bunting population aligns with the pattern of isolation by distance (IBD), we conducted a correlation analysis between their geographic distribution and genetic distance. We calculated the genetic distance (*Dij*) between each pair of samples using RapidNJ v2.3.2 (Simonsen et al., 2008), while R package geosphere was used to calculate the geographic distance (Hijmans et al., 2017). To test the statistical significance in isolation by distance analysis, we carried out the Mantel test using R package vegan (Dixon, 2003). In addition, we performed isolation by distance analysis for each pair of populations. The genetic distance between populations was determined using pairwise *F*_ST_. The geographical coordinates of the central point of each population were used to calculate the geographic distance between them.

### 2.6 Estimation of gene flow

Three methods were used to estimate gene flow between different populations of yellow-breasted bunting. We calculated the Patterson’s *D* statistic (also called ABBA-BABA statistics) for each trio of yellow-breasted bunting populations using the script developed by Simon Martin (https://github.com/simonhmartin/genomics_general). The setting of each trio is described in Table S3. The significant test, the block jackknife procedure, was applied to the mean genome-wide *D* values, with |Z| score >3 indicating the *D* values deviate significantly from zero (Patterson et al., 2012). The significant deviation of *D* values suggests the presence of gene flow between populations A and C or populations B and C (Table S3).

The second method involved estimating the *f-branch* (*f_b_*) metric using autosomal SNPs with Dsuite (Malinsky et al., 2021). This approach extends from Patterson’s *D* statistic, further distinguishes corrected f4-ratio results, and assigns gene flow evidence to specific branches on a phylogeny. The phylogeny used in this analysis was obtained from the ML tree of autosomal SNPs, where reed bunting and black-faced bunting were the outgroups.

Individuals with uncertain phylogenetic relationships or hybrids of different populations were excluded from this analysis. The significance of each *f_b_* ratio was evaluated using the Z-scores and its associated *p*-values. The third method involved estimating the number and direction of past migration events using Treemix (Pickrell & Pritchard, 2012) based on autosomal SNPs. To determine the appropriate number of migration edges (m) on the population tree, we inferred the optimal value of m using OptM (Fitak, 2021). We ran 100 bootstrap replicates for each edge (from 1 to 10) with 1000 SNP blocks to account for the LD and find the best m. Finally, we used the matrix of residuals to establish metrics for evaluating the model’s goodness of fit to the data.

### 2.7 Inference of demographic history

We used three methods to reconstruct the demographic history from ancient to recent times. We first inferred the changes in past effective population size from individual whole-genome sequences using pairwise sequentially Markovian coalescent (PSMC) (Li & Durbin, 2011). To avoid bias from low coverage and high missing data, we calculated each individual’s sequencing coverage and missing rate. These individuals had an average sequencing depth of 24.6×, with the lowest sequencing depth being 16.3×. The average missing rate was 0.46%, and only one individual had a missing rate greater than 10% (i.e. 13.2%). Therefore, all individuals were used for PSMC analysis. We mapped the clean reads of each individual to the reference genome using BWA and then called the SNPs using SAMtools. The analysis excluded sex-linked SNPs. The PSMC was carried out with the default setting (-N25 -t15 -r5 -p “4+25*2+4+6”), and the mutation rate was set to 6.9×10^−9^ per site per generation (2.3×10^−9^ per site per year) (Smeds et al., 2016). The mutation rate came from the collared flycatcher (*Ficedula albicollis*), as this data was not available for the yellow-breasted bunting. Based on previous studies, we set the generation time to three years (BirdLife International, 2024).

We also inferred recent demographic history for each yellow-breasted bunting population using SMC++ (Terhorst et al., 2017). SMC++ uses a new spline regularization method that combines site frequency spectrum (SFS) and LD information in coalescent HMMs to improve estimation accuracy (Terhorst et al., 2017). To avoid bias caused by selection and other factors, we excluded SNPs in the sex chromosome and coding regions. We performed SMC++ analysis with the following parameters: --polarization-error 0.5 --spline piecewise -- regularization-penalty 6.0 --knots 20. The mutation rate used in this analysis was 6.9×10^−9^ per site per generation, and the generation time was set to three years.

We estimated the effective population size of the past 700 generations using GONE (Santiago et al., 2020). This analysis was based on the observed spectrum of linkage disequilibrium (LD) of pairs of loci. We excluded the SNPs in the sex chromosome and coding regions. Due to the limitation of the software on the number of SNPs on each chromosome, we used scaffold-level data instead of pseudochromosome-level data for this analysis. Specifically, we used 191 scaffolds that were longer than 1Mb, and the number of SNPs analyzed in each scaffold was 50,000. We repeated the analysis 200 times for each population. We used the recombination rate (1.5cM/Mb) of the zebra finch (*Taeniopygia guttata*) (Backstrom et al., 2010) in this analysis.

### 2.8 Ecological niche modeling

To predict changes in species niches and distribution in past and future generations, we conducted ecological niche modeling (ENMs) analysis using Maxent (Phillips et al., 2006). This method applies a machine-learning technique called maximum entropy modeling to predict the most suitable conditions for a species based on environmental data and occurrence records. We obtained occurrence records from GBIF (https://www.gbif.org/species/2491518) and eBird (https://ebird.org/species/yebbun). We filtered the records based on the following criteria: for the prediction of the breeding habitat, we only used records from the breeding season (June to August) and within the breeding area; for the prediction of the wintering area, we used records from the wintering season (November to February) and within the wintering area. Then, we performed spatial thinning (keeping only one record within a range of 1 km^2^). Then, we downloaded 19 bioclimatic variables from WorldClim (www.worldclim.org), but we excluded 13 bioclimatic variables from the analysis based on correlation analysis and permutation importance analysis. Finally, we predicted the suitable breeding and wintering ranges of yellow-breasted bunting at the Last Interglacial (LIG), Last Glacial Maximum (LGM), Mid-Holocene (MH), the current day, and the year 2070. For the ENMs analysis of the year 2070, we ran the analysis under two Representative Concentration Pathways (RCP8.5 and RCP2.6) (van Vuuren et al., 2011), which represent the highest and lowest of the four greenhouse gas concentration pathways.

### 2.9 Genetic diversity and inbreeding analyses

To better understand the long-term evolutionary potential of yellow-breasted bunting, we estimated the genome-wide genetic diversity. We calculated the nucleotide diversity (π) based on 20kb non-overlapping sliding windows using VCFtools 0.1.1421. Additionally, we estimated the genome-wide heterozygosity for each individual by dividing the total number of heterozygous sites by the effective length of the genome.

We estimated the inbreeding level by identifying the Runs of Homozygosity (ROH) using PLINK v.1.919. The analysis was performed with the following settings: --mendel –genome --homozyg --homozyg-group --homozyg-window-snp 50 --homozyg-snp 50 --homozyg-window-missing 3 --homozyg-kb 100 --homozyg-density 50 --homozyg-window-het 3. We categorized the ROH into two types based on their length: short-ROH (0.1 Mb < ROH < 1Mb) and long-ROH (ROH > = 1Mb) (Ceballos et al., 2018). We defined the genome-wide inbreeding coefficient, *F_ROH_*, as the total length of ROH divided by the effective length of the genome. Additionally, we estimated the distribution of ROH by calculating the *S_ROH_* (total length of ROH) and *N_ROH_* (total number of ROH).

### 2.10 The accumulation of deleterious mutations

To assess the genetic load in different populations of yellow-breasted bunting, we counted the number of deleterious mutations present in the genome. We first defined the derived alleles using the reed bunting as an outgroup. If an allele is present in the yellow-breasted bunting population and has a zero frequency in the outgroup, it is classified as a derived allele. Then, we used snpEff v.4.3 (Cingolani et al., 2012) to annotate the functional effect of derived alleles, categorizing them as loss-of-function (LoF) variants, nonsynonymous variants, and synonymous variants. LoF variants are considered the most harmful mutations and contain variations with splice donor, splice acceptor, start lost, stop lost, stop gained, and stop retained mutations. Nonsynonymous mutations were further classified into two groups based on Grantham’s score (Grantham, 1974): deleterious (Grantham’s score > 150) and tolerated (Grantham’s score =< 150) nonsynonymous mutations. The synonymous variants were considered neutral.

We calculated the total number of four types of mutations - LoF, deleterious nonsynonymous, tolerated nonsynonymous, and synonymous mutations - for each individual. We also calculated the number of homozygotes and heterozygotes for these four types of mutations. Additionally, we used the ratio of deleterious mutations to neutral mutations as the index of effectiveness of purifying selection (Robinson et al., 2022). We calculated the ratio of LoF, deleterious nonsynonymous, and tolerated nonsynonymous mutations for each population of yellow-breasted bunting.

### 2.11 Simulations

To evaluate the impact of recent demographic history on the accumulation of deleterious mutations, we performed forward simulations with the Wright-Fisher (WF) model using SLiM v. 4.0.1 (Haller & Messer, 2023). We simulated the most recent 500 generations of demographic history obtained from each population’s GONE result. To reduce computational time, we scaled down the effective population size of each population to 0.02-fold and simulated a genome with 14000 genes for each individual. Each gene has a length of 1650 bp. These genes were distributed across 26 chromosomes. The mutation rate was set to 4.6×10^-9^, and the recombination rate within genes was set to 1×10^-8^. We assumed no recombination between genes, while the recombination between chromosomes is free. The proportion of deleterious mutations was defined based on the distribution of fitness effect (DFE), which was calculated using polyDFE v2.0 (Tataru & Bataillon, 2019). We modified the “hmix” model based on the previous studies (Kyriazis et al., 2023; Kyriazis et al., 2021) for the setting of dominance coefficients.

To conduct our simulations, we first ran a burn-in period of 10 times the effective population size. During this period, we sampled the data every 1000 generations. After the burn-in period, we sampled the data every two generations. For data collection, we obtained data from a sample of 40 individuals from the population. The data we collected during the simulation included four types of mutations: strongly deleterious (s ≤ −0.01), moderately deleterious (−0.01 < s ≤ −0.001), weakly deleterious (−0.001 < s ≤ −0.00001), and neutral alleles (s=0.0). We conducted 30 replicates for each simulation.

## 3 Results

### 3.1 Genome assembly and evaluation

We sequenced and assembled a high-quality reference genome of yellow-breasted bunting using the 10× genomics sequencing method. The reference genome size is 1.13 Gb, with a scaffold N50 of 4.57 Mb and a contig N50 of 115 kb. BUSCO analysis showed that the reference genome had 91.5% completeness (Table S2). We annotated 16671 protein-coding genes for the reference genome. We performed whole-genome resequencing on 94 yellow-breasted bunting individuals and two outgroups (Figure 1b). The mean sequencing depth was 24.6×, and the mean missing rate was 0.46%. Lastly, we identified 145,174,637 autosomal biallelic SNPs and 1,319,040 Z-linked SNPs for downstream analyses.

### 3.2 Population structure and gene flow

We used three methods to estimate the population structure and genetic divergence within this species. The ML tree based on autosomal SNPs formed three main populations: populations P1 and P3 were monophyletic clades, while P2 was paraphyletic to P1 and P3 (Figure 1a). P1 was further divided into two monophyletic subpopulations, P1.a and P1.b. A similar pattern was also observed in the ML tree based on Z-linked SNPs, although the clusters of P2 were unstable (Figure S1). The PCA results based on autosomal and Z chromosomal SNPs (Figure 1c and 1d) were consistent with the phylogenetic tree: three main clusters were formed, with P1 divided into two clusters on PC2 and P2 in the middle of P1 and P3. The result of Admixture mainly supports the populations of P1 and P3, while the population P2 appears to be a mixture of P1 and P3 (Figure 1e). K=1 is the most supported number of clusters based on the result of CV error. However, the optimized K can be unreliable, especially when the population differentiation is shallow (Kalinowski, 2011). Moreover, according to the ML tree results, P2 did not form a monophyletic clade with the other two populations. We therefore consider P2 to be a different population and hypothesize that it could be the parental population (see Discussion). Both *F*_ST_ and the mitochondrial genome results suggest that the genetic differentiation between these populations was very shallow. The average genome-wide *F*_ST_ between these populations is lower than 0.02, with the highest *F*_ST_ between P1 and P3, particularly P1.b and P3 (*F*_ST_ =0.017), and the lowest *F*_ST_ between P2 and P3 (*F*_ST_ =0.007) (Figure 2a). Moreover, no clear structure was found in the PCA and ML tree based on the mitochondrial genomes (Figure S2). This could be due to the shallow differentiation between different populations and the relatively small amount of genetic information provided by mitochondria.

**Figure 2.**
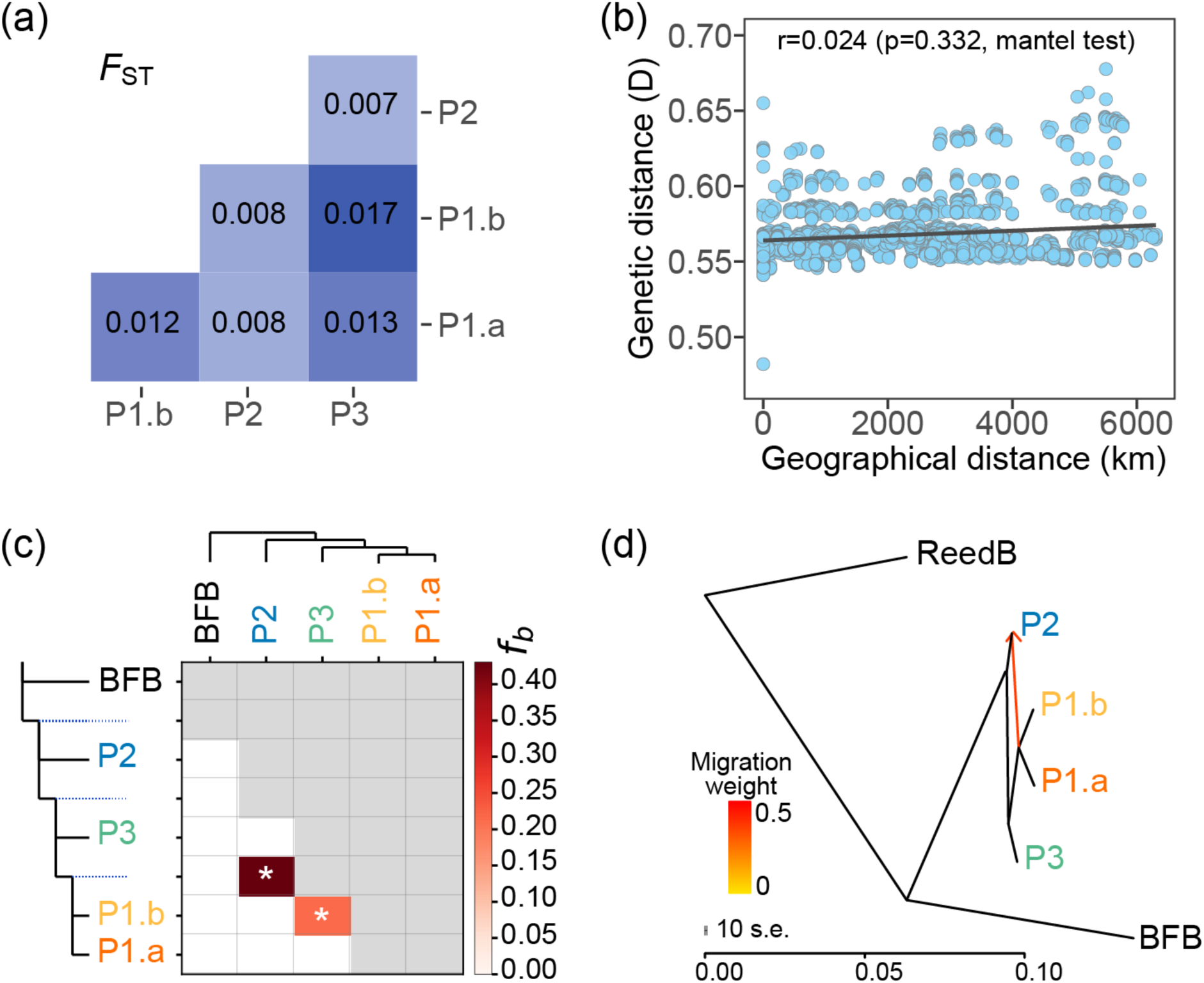
The genetic differentiation and gene flow between different populations of yellow-breasted bunting. (a) The matrix of genetic differentiation, *F*_ST_, between different groups. The color indicates the degree of genetic differentiation. (b) The correlation between the genetic distance and the geographic distance. (c) The estimated gene flow using branch-specific statistic method (*f_b_*) with Dsuite. The value in the matrix (*f_b_*) indicates the excess allele sharing between populations. The asterisks (*) indicate significant *f_b_* value (p<0.01). (d) The estimated number and direction of migration events using Treemix. The color of the arrows is based on the migration weight.

The populations of yellow-breasted buntings were associated with the geographical distribution (Figure 1b). P1 was located in the eastern regions, including the island population (P1.a: Sakhalin, Japan, and Kuril Islands) and coastal areas (P1.b: Primorskiy). P2 was located in the North-Eastern Asia area, between P1 and P3. P3 occupied most of the distribution range from Finland in Europe, Mongolia, to Eastern Siberia. Although these populations were highly correlated with geographic distribution range, they did not show an “isolation by distance” relationship based on genetic distance between individuals (Figure 2b) nor the pairwise *F*_ST_ between populations (Figure S4).

Three methods were applied to investigate whether the weak differentiation among these populations was caused by gene flow. Patterson’s *D* statistics revealed significant gene flow between populations P2 and P3, as well as gene flow between P2 and P1.a, and gene flow between P3 and P1.a (Table S3). The result of the *f-branch* (*f_b_*) metric identified two significant gene flows between these populations (Figure 2c). Most of the migration events occurred between P2 and P1 (*f_b_* = 43.04%), and the migration events between P3 and P1.b amounted to 21.55%. Treemix results indicate one migration edge (m) on the population tree, suggesting that gene flow happened between P2 and P1 (Figure 2d, Figure S3).

### 3.3 Demographic history and climate change

The PSMC analysis based on individual genomes supported similar population fluctuations of the three populations in ancient history (Figures 3a-c). All populations showed an effective population expansion before the Last Interglacial (LIG), followed by different degrees of population decline. The population size of P1 continued to decline during the last glacial period (LGP), while the population size of P3 showed a similar decline but rebounded slightly during LGP. In contrast, the population size of P2 did not show a significant decrease even after the LIG.

**Figure 3.**
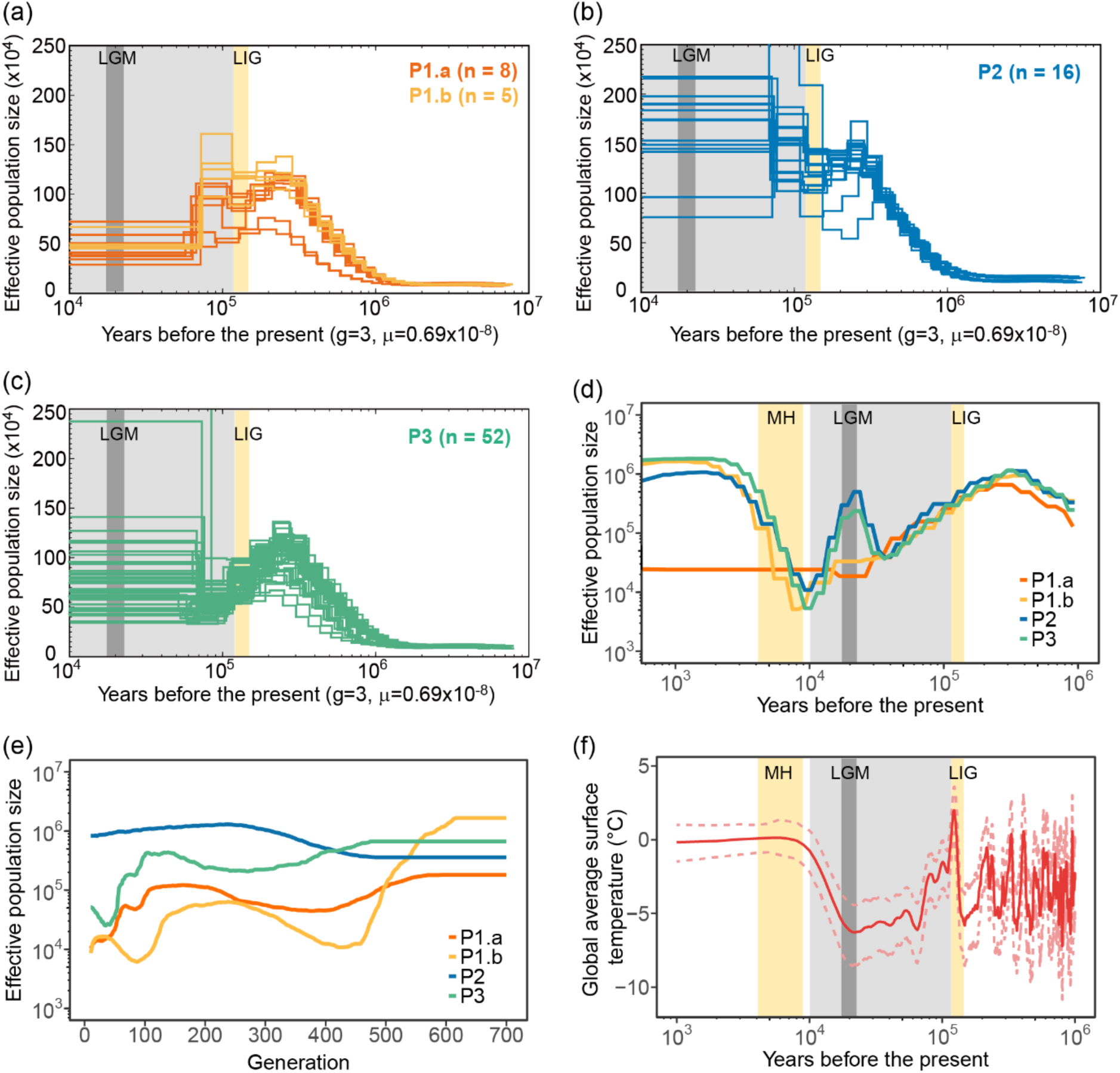
Demographic histories of yellow-breasted buntings. (a) Demographic history of the population (a) P1.a and P1.b, (b) P2, and (c) P3 based on the individual autosomal SNPs using pairwise sequentially Markovian coalescent (PSMC). (d) The inference of effective population size using SMC++. (e) The estimation of effective population size in the past 700 generations using GONE. (f) The changes in global average surface temperature in the past millions of years. The yellow color blocks indicate the Last Interglacial (LIG) and Mid-Holocene interglacial periods, respectively. The light gray color block indicates the Last Glacial Period (LGP), while the dark grey color block indicates the Last Glacial Maximum (LGM).

The relatively recent demographic history inferred using SMC++ suggested that these populations shared similar demographic trajectories before the Last Glacial Maximum (LGM) but showed different responses to LGM (Figure 3d). The population size of P1.a had been stable but relatively low since the LGM, while the population size of P1.b showed an expansion during the mid-Holocene (MH) period after a long-term decline. On the other hand, the population fluctuation of P2 and P3 was similar. Both populations rebounded before LGM and declined again since the LGM, and they increased in population size again after entering MH.

We further inferred the changes in the population size over the past 700 generations (∼2,100 years) using GONE. P1.a, P1.b, and P3 experienced similar population trajectories, all undergoing a bottleneck during the 300-500 generations and a population decline in the past 100 generations (Figure 3e). The effective population size of P1.a and P1.b was lower than that of P3. However, the population size of P2 remained high and stable over the past 700 generations. Therefore, P1.a and P1.b experienced a more severe recent population decline than the other two populations.

The population fluctuation of these populations appears to be highly related to climate change. The changes in global average surface temperature (Snyder, 2016) were consistent with the fluctuation pattern of effective population size (Figure 3f). During most of the LGP, with lower temperatures, most populations experienced a population decline. In contrast, most populations except P1.a underwent a population expansion during the warmer MH period. Moreover, climate change also impacted the suitable breeding area (Figure 4). The ENM results indicated that the breeding range of the yellow-breasted bunting underwent a contraction during the LIG period, as compared to its existing distribution area. The suitable breeding range reached a minimum during the LGM period, with an overall shift towards the south. The breeding habitat showed an east-west discontinuity during the LIG and LGM periods. However, during the MH period, suitable habitats increased with the rise in temperature. Unlike the breeding range, the wintering area did not show obvious differences in the ancient period (Figure S5).

**Figure 4.**
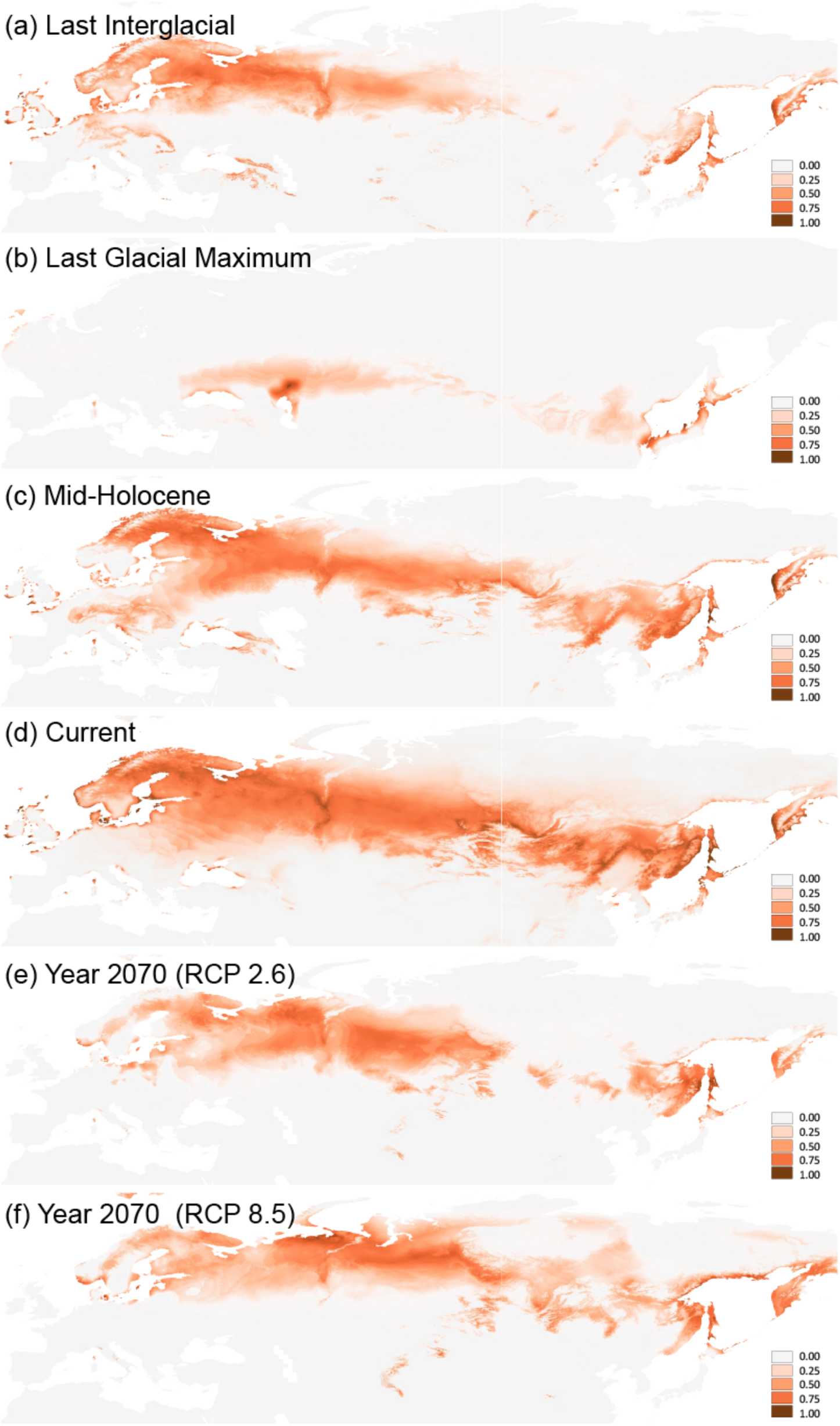
The predicted suitable breeding range of yellow-breasted bunting. Reconstruction of ecological niche models for the breeding area of yellow-breasted buntings in the Last Interglacial (LIG), Last Glacial Maximum (LGM), Mid-Holocene (MH), current, and year 2070 with different Representative Concentration Pathways (RCP2.6 and RCP 8.5). The color indicates the probability of suitable conditions in the models (i.e. darker color indicates more suitable).

To investigate the impact of climate change on the yellow-breasted bunting in future generations, we further predicted the suitable breeding and wintering area of this species in year 2070. We conducted the ENM analysis based on different climate change scenarios, specifically, the Representative Concentration Pathways (RCP) 2.6 and 8.5. Both ENM results suggest that there will be a contraction in the suitable breeding area in the year 2070, especially in the middle regions (i.e., Mongolia and NE Asia) and the eastern regions (i.e., coast and islands) (Figure 4e-f). In contrast, there are no significant impacts on the wintering area in year 2070 (Figure S4e-f).

### 3.4 Genetic diversity and inbreeding level

Genetic diversity is a critical factor that reflects the long-term evolutionary potential of a species, which could be affected by demographic history and inbreeding. The mean nucleotide diversity of yellow-breasted bunting was 5.67×10^-3^, with specific nucleotide diversity for P1.a, P1.b, P2, and P3 was 5.44×10^-3^, 5.78×10^-3^, 6.00×10^-3^, and 5.57×10^-3^, respectively. The overall genome-wide heterozygosity was 4.88×10^-3^, with specific heterozygosity for P1.a, P1.b, P2, and P3 was 4.67×10^-3^, 4.87×10^-3^, 5.13×10^-3^, and 4.84×10^-3^, respectively. Our results indicate that P1.a has a significantly lower nucleotide diversity and genome-wide heterozygosity than other populations (Figure 5a-b). This might be due to the long-term population decline and the relatively low effective population size of this population. On the other hand, P2, which experienced a large and stable population size, has higher nucleotide diversity and genome-wide heterozygosity than other populations. The inbreeding levels varied among different populations of yellow-breasted buntings.

**Figure 5.**
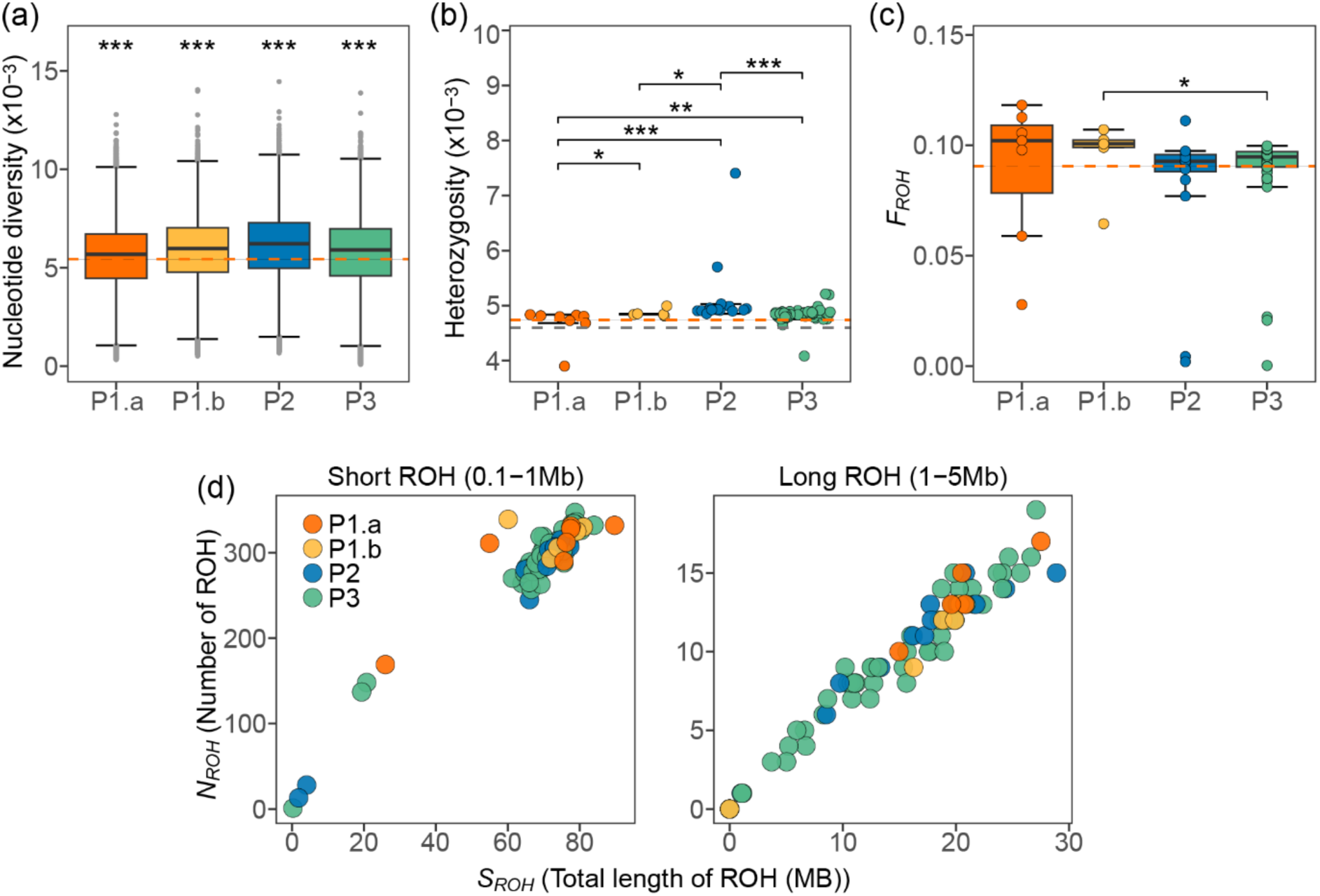
The genome-wide genetic diversity and level of inbreeding of yellow-breasted buntings. The estimated (a) average nucleotide diversity and (b) genome-wide heterozygosity of different populations of yellow-breasted buntings. (c) The genome-wide inbreeding coefficient, *F_ROH_*, of different populations. ROHs longer than 100kb were included in this analysis. The red dotted lines in (a), (b), and (c) indicate the mean nucleotide diversity, mean genome-wide heterozygosity, and mean *F_ROH_* of population P1.a. The grey dotted line in (b) indicates the mean genome-wide heterozygosity (i.e. 4.6×10^-3^) of yellow-breasted buntings collected in 2019 based on Wang et al. (2022). (d) The distribution of the total number of ROHs and the total length of ROHs is shown for two categories of ROHs. Short-ROH (left): ranges from 0.1 to 1 Mb; long-ROH (right): longer than 1 Mb. The ‘***’ in (a) indicates that the differences between all population pair are significant. In (b) and (c), significant differences between population pairs are indicated by asterisks (*). * p<0.05; ** p<0.01; *** p<0.001.

Populations P1.a and P1.b have a higher *F_ROH_* compared to populations P2 and P3, although only the difference between P1.b and P3 is significant (Figure 5c). This trend is also observed in short-ROH and long-ROH (Figure S6). There was no clear difference in the *N_ROH_* and *S_ROH_* among individuals from different populations (Figure 5d). This implies that these populations might have undergone similar demographic histories in both recent (long-ROH) and relatively ancient (short-ROH) times.

### 3.5 The accumulation of deleterious mutations

Identifying and measuring deleterious mutations is vital for the conservation management of endangered species since they can help us assess the extinction risk of a species. We found that P1.a and P1.b had accumulated more homozygous deleterious mutations compared to P2 and P3 (Figure 6a). This pattern was observed in both highly deleterious mutations (LoF), deleterious nonsynonymous, and tolerated nonsynonymous mutations. However, the difference between the total number of deleterious mutations and heterozygous deleterious mutations was insignificant among these populations (Figure S7). On the other hand, population P2 has the lowest number of homozygous LoF, deleterious nonsynonymous, and tolerated nonsynonymous mutations.

**Figure 6.**
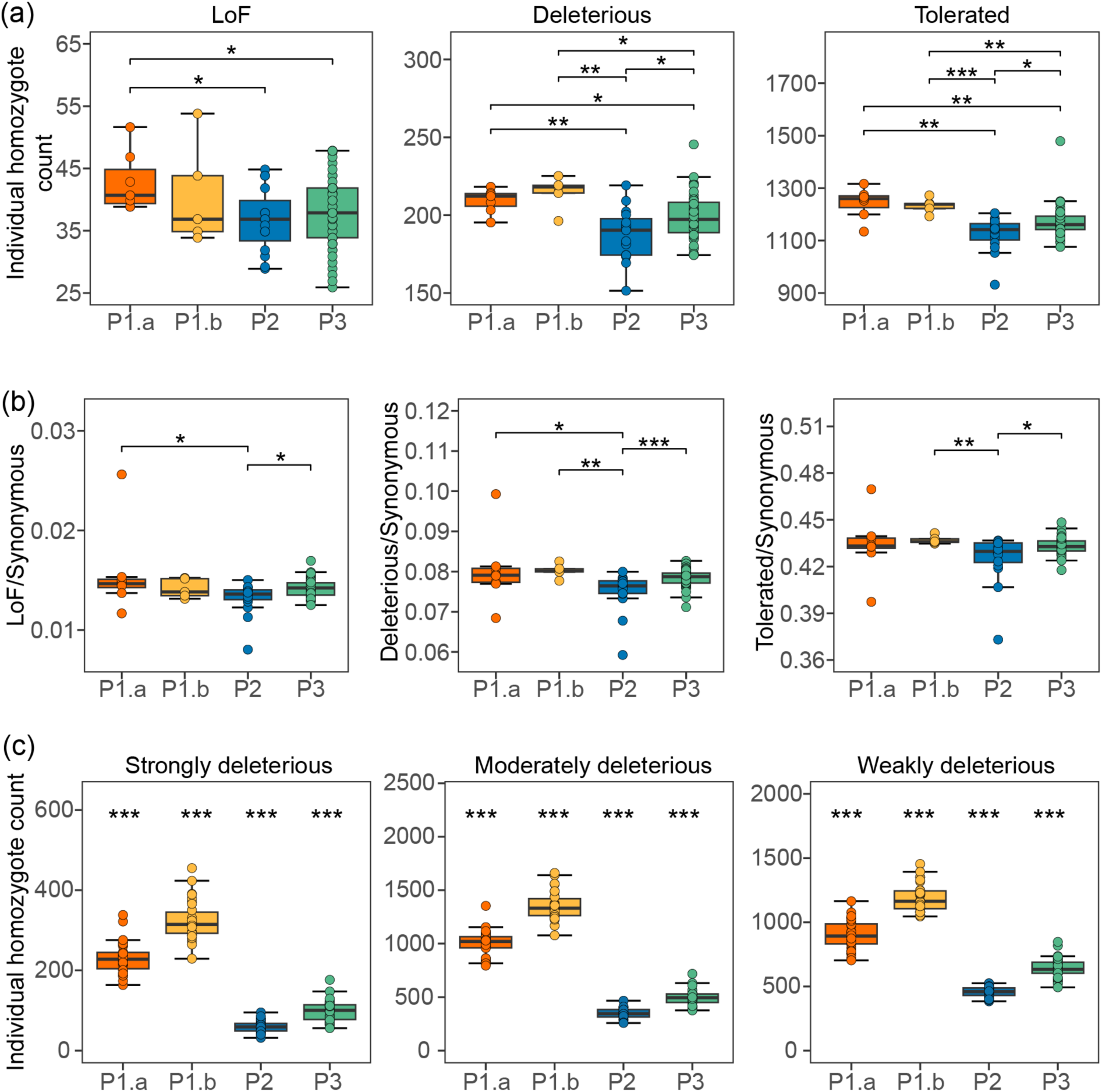
The accumulation of deleterious mutations in the yellow-breasted bunting populations. (a) The number of homozygous loss-of-function (LoF), deleterious nonsynonymous, and tolerated nonsynonymous mutations per individual. (b) The ratio of LoF, deleterious nonsynonymous, and tolerated nonsynonymous mutations to synonymous mutations. (c) The number of homozygous strongly, moderately, and weakly deleterious mutations per individual based on the simulations of the demographic history of the yellow-breasted bunting populations. We simulated the demographic scenarios of the past 500 generations of yellow-breasted buntings. In (a) and (b), the significant differences between population pairs are indicated by asterisks (*). The *** in (c) indicates a significant difference between population pairs. * p<0.05; ** p<0.01. *** p<0.001.

We evaluated the effectiveness of purifying selection using the ratio of deleterious mutations to neutral mutations. The effectiveness of selection is associated with the effective population size, with larger populations generally experiencing stronger selection. Our results indicate that the differences in the ratio of deleterious mutations are insignificant among most populations, except that P2 has a significantly lower ratio of LoF, deleterious and tolerated nonsynonymous mutations (Figure 6b).

To investigate whether the accumulation of homozygous deleterious mutations in P1 is due to demographic changes, we simulated the demographic scenarios over the past 500 generations for the yellow-breasted bunting populations. The simulation results indicate that P1.a and P1.b have accumulated more homozygous deleterious mutations (including strongly, moderately, and weakly deleterious mutations) than P2 and P3 under corresponding demographic scenarios (Figure 6c). In addition, population P2 has accumulated fewer homozygous deleterious mutations than other populations.

## 4 Discussion

### 4.1 Shallow genetic differentiation among yellow-breasted bunting populations

We have identified three populations of the yellow-breasted bunting with shallow genetic differentiation between them. Our findings suggest that the genetic divergence among these populations is primarily associated with their geographic distribution, spanning from east to west. The genetic differentiation between these populations was notably low (*F*_ST_ < 0.02), possibly due to gene flow between them. All three methods have uncovered gene flow between P2 and P1, and some methods have identified gene flow between P3 and P1.a or P1.b, as well as between P3 and P2. The inconsistency in the results of these methods could be due to method sensitivity and the strength of gene flow. Previous studies suggest the strength of gene flow, the direction of gene flow, the number of SNPs, and the effective population size can impact the accuracy of *D* statistics and *f*-branch statistics (Malinsky et al., 2021; Patterson et al., 2012; Zheng & Janke, 2018). These methods may not detect every gene flow event and are most robust when gene flow is particularly strong (Malinsky et al., 2021). Prior research indicates that the uneven gene flow between populations may be attributed to geographic distances within species (Wang, 2013; Wang et al., 2013). However, the genetic distance of the yellow-breasted bunting does not follow the pattern of isolation by distance (IBD). This could be attributed to P3 occupying the majority of the distribution area, with minimal differentiation within P3, while the primary genetic divergence occurs between P1 and P3, leading to the absence of IBD.

Based on the population structure and geographic distribution of these populations, we have formulated two hypotheses that could explain the population structure of the yellow-breasted bunting. First, population P2, situated in the middle, may represent an admixed population of P1 and P3. Second, population P2 may serve as the parental population of yellow-breasted buntings, with P1 and P3 expanding to the east and west from P2, respectively. This hypothesis is predominantly based on the autosomal phylogenetic tree, which situates P2 at the tree’s root. Furthermore, P2 had a larger and more stable effective population compared to the other two populations, suggesting its role as the parental population. Although both hypotheses predict a higher genetic diversity and lower *F*_ST_ in P2, the position of P2 on the phylogenetic tree suggests the second hypothesis to be a more probable scenario. However, the two hypotheses are not mutually exclusive and might have both contributed to the genetic structure of the yellow-breasted bunting.

These identified populations partly correspond to the distribution of subspecies of yellow-breasted buntings (Copete & Sharpe, 2020; Park et al., 2020). P3 falls entirely within the range of subspecies *E. a. aureola*, while P1 falls entirely within the range of *E. a. ornata* (Figure 1b). The main disagreement occurred in the East Transbaikalia, North-Eastern China, Anadyr land, and Kamchatka. Based on the genetic structure, the population in Anadyr land and Kamchatka (P3.a, highlighted in black) should belong to P3, and birds in this region likely spread from the middle region of P3. The population in East Transbaikalia also belongs to P3. On the other hand, birds in the North-Eastern China area should belong to P2.

### 4.2 The impacts of climate change and human activities on ancient and recent demographic history

Most populations of this species showed a similar pattern but different degrees of decline from the past thousands to millions of years. The demographic patterns and effective population size of PSMC results of populations P1 and P3 were generally consistent with the previous genomics study on yellow-breasted bunting (Wang et al., 2022), which did not identify the demographic history pattern of P2. For the relatively recent demographic history, we have identified two bottlenecks in populations P2 and P3 during the LGP using SMC++, while Wang et al.’s (2022) study failed to reconstruct the demographic trajectories during this period. This may be due to the Stairwayplot having a lower resolution in recent history when the sample size was small (Liu & Fu, 2015). Furthermore, our results suggest that the ancient demographic history of yellow-breasted bunting was mainly affected by global climate change. Specifically, the population of yellow-breasted bunting experienced significant declines during the LGP while showing a notable population expansion during the warmer MH period. Additionally, the changes in the suitable breeding range of yellow-breasted bunting were also in line with the population fluctuations, with the minimum distribution range occurring during the LGM. This implies that the shrinking habitat forces the populations to contract towards glacial refuges during the LGM. The combination of harsh environments and fragmented habitats resulted in population declines during the LGM (Hewitt, 2004; Nadachowska-Brzyska et al., 2015). Our study aligns with previous research on 38 avian species, suggesting that many bird species experienced population contractions and expansions during the Quaternary period, coinciding with climate cycles (Nadachowska-Brzyska et al., 2015).

However, population fluctuations during the glacial period can vary among populations, even within the same species. For instance, unlike populations P2 and P3, the P1 population remained at a low level of effective population size after a long-term decline during the LGP (Figure 3d). Subsequently, the P1.b population recovered during the MH, while P1.a did not. This difference may be related to the geographic location of the populations. P1.a primarily inhabits islands, which are more susceptible to the effects of climate change. The cooler and drier climate during the LGM could lead to changes in vegetation zones and the fragmentation and contraction of island species’ habitats (Collins et al., 2013; Tsukada, 1983). Additionally, the drought may have led to a decline in food resources (such as seeds and insects) for the species, potentially contributing to a decrease in populations (Albright et al., 2010; Tsukada, 1983).

Human activities have likely influenced the recent demographic history of the yellow-breasted bunting, as well as other endangered avian species (Dierickx et al., 2020). Over the past 10,000 years, we were in the stable and warm “Holocene” interglacial period. However, the populations of yellow-breasted bunting have undergone varying demographic trajectories. During the past 700 generations, Population P2 remained large and stable, while both P1 and P3 encountered two bottlenecks. The first bottleneck, which occurred 400-500 generations ago, primarily affected the restricted coastal population P1.b, leading to a drastic 100-fold decline in population size. The second bottleneck, around 100 generations ago (approximately 300 years ago), had a broader impact, resulting in a 10-fold population decline for both P1 and P3. We speculate that the recent decline in the population of yellow-breasted bunting may be attributed to human activities, such as the Second Agricultural Revolution in Europe, spanning from the mid-17^th^ to late 19^th^ centuries (Mingay, 1977). This revolution accelerated land development and standardization, along with the widespread use of fertilizers (Mingay, 1977; Thompson, 1968). Previous researches suggest that agricultural intensification, especially increased use of pesticides and chemical fertilizers, likely reduced food availability for many farmland bird species by diminishing insect and invertebrate populations, particularly during breeding seasons (Donald et al., 2001; Rigal et al., 2023).

Consequently, the agricultural intensification in Europe and other regions may have impacted the population size of the yellow-breasted bunting in its breeding area over the past 100 generations. Furthermore, subsequent human trapping during migration has exacerbated the decline in the population of this species (Kamp et al., 2015).

### 4.3 The impact of demographic histories on genetic features

The genetic diversity of yellow-breasted buntings was strongly associated with their demographic history. Populations P1 and P3, which have undergone two recent bottlenecks, exhibit significantly lower levels of genetic diversity compared to P2. In particular, population P1.a, which has remained a smaller effective population size since LGM, shows the lowest genetic diversity. Additionally, the demographic history has also influenced the level of inbreeding in this species. Two bottlenecks over the past 700 generations have likely resulted in higher inbreeding in populations P1 and P3 compared to P2, although this difference was not significant. Moreover, the smaller effective population size has probably led to a higher level of inbreeding in population P1.b than in population P3, even though they have similar demographic trajectories. The demographic history of populations also had a significant impact on the accumulation of deleterious mutations. While these populations have similar total deleterious mutations, populations P1.a and P1.b have significantly more homozygous deleterious mutations compared to the other two populations. The relatively small effective population size and high inbreeding levels in populations P1.a and P1.b may contribute to the accumulation of homozygous deleterious mutations. This phenomenon has also been observed in species such as Indian tigers and Alpine ibex (Grossen et al., 2020; Khan et al., 2021).

Simulations further support the idea that populations having lower effective population sizes accumulate relatively more homozygous deleterious mutations. Furthermore, we found that differences in homozygous deleterious mutations among populations in the simulation results were more pronounced than those in the empirical data. This may be due to gene flow between populations in real scenarios, which may affect the accumulation of deleterious mutations in different populations (Couvet, 2002). However, unlike other endangered species that have gone through long periods of small population sizes (Grossen et al., 2020; Kardos et al., 2023), the three yellow-breasted bunting populations still have relatively large effective population sizes, suggesting stronger purifying selection overall, like the passenger pigeon (Murray et al., 2017). However, there were differences in the strength of purifying selection among the populations. We did not find a significant difference in the ratio of deleterious mutations between populations P1 and P3, which may be due to their similar effective population sizes. In contrast, population P2 had a significantly lower ratio of deleterious mutations compared to the other populations, possibly due to the larger effective population size, resulting in stronger purifying selection in this population. This is in line with previous studies suggesting that the ratio of deleterious mutations is negatively correlated to effective population sizes, implying stronger purifying selection in larger populations (Bertorelle et al., 2022; Robinson et al., 2022).

Although the yellow-breasted bunting experienced population fluctuations similar to those of the passenger pigeons during the LGP, it did not undergo regular fluctuations as the passenger pigeons did before the LGP (Hung et al., 2014). In addition, PSMC results show that the historical *N_e_* of yellow-breasted bunting ranged from 10^5^ to 2×10^6^, which is much larger than that of passenger pigeons (ranging from 4×10^4^ to 2×10^5^). Meanwhile, the yellow-breasted bunting exhibited a genome-wide nucleotide diversity (5.67×10^-3^) that is twice that of the passenger pigeon (π = 2.7×10^-3^). Moreover, the yellow-breasted bunting’s genome-wide heterozygosity (4.88×10^-3^) is much higher than that of other endangered bird species (ranging from 0.43×10^-3^ to 1.88×10^-3^) (Dussex et al., 2021; Li et al., 2014; Zhan et al., 2013). However, the yellow-breasted bunting remains at risk of extinction, particularly in light of recent population collapse. The case of the passenger pigeons serves as a stark reminder that even species with larger populations and the capacity to purge harmful mutations can still face extinction when confronted with significant environmental changes (Murray et al., 2017). Furthermore, larger populations generally contain more deleterious mutations, rendering the species more prone to inbreeding depression during substantial population declines (Bertorelle et al., 2022). Moreover, the recent population collapse likely exacerbated the genetic threats in the current population, leading to an overestimation of this species’ genetic health. Indeed, when we compared the average heterozygosity estimated in the current study (4.88×10^-3^; based on samples collected from 1992 to 2004) with that from an earlier study based on a small number of samples collected in 2019 (4.6×10^-3^; Wang et al., 2022), the more recent samples had an average heterozygosity lower than that of P1.a and much lower than that of the other populations. This suggests a possible genetic diversity decline in the yellow-breasted bunting within the last 30 years.

### 4.4 Conservation implications

Our findings indicate the necessity to prevent the loss of genetic diversity and the increase of inbreeding in the yellow-breasted bunting. While the population before the recent population collapse exhibited relatively low genetic threats with high genetic diversity and low levels of inbreeding, the genetic threats faced by the current population remain unclear. It is essential to take measures to halt the ongoing decline of the population and to conduct regular quantitative and genetic monitoring. The yellow-breasted bunting population is primarily threatened by illegal hunting along migratory routes. Although China has prohibited the trade of this species since 1997 and has made improvements in law enforcement and awareness campaigns in recent years (Heim et al., 2021; Kamp et al., 2015), more attention is still required to address this issue. The other significant threat is the impact of climate change and habitat loss. Our results suggest that although climate change will not significantly affect the wintering area by 2070, it will lead to a significant reduction in the breeding area. The breeding grounds in the eastern distribution range will largely disappear, and the future breeding area will contract to North-Eastern Europe and Western Siberia. Farmland birds in Europe are facing ongoing population declines due to agricultural intensification (Donald et al., 2001; Rigal et al., 2023). Additionally, migratory birds such as the yellow-breasted bunting may be more vulnerable to climate change due to their limited potential for range shifts in comparison to resident birds, since they exhibit higher fidelity to breeding and wintering areas (Välimäki et al., 2016). Therefore, it is crucial to safeguard the habitat and food resources within the breeding range of the yellow-breasted bunting to mitigate the impact of climate and environmental changes.

In light of the genetic structure findings, we propose conserving the yellow-breasted bunting as one conservation unit, given the shallow genetic differentiation among the three populations and the gene flow between them. However, additional conservation efforts are necessary for the P1 population, which consistently exhibits a lower effective population size and genetic diversity, along with more homozygous deleterious mutations. If these exposed deleterious mutations are not promptly purged, they could impact the population’s overall fitness (Hedrick & Garcia-Dorado, 2016). Specifically, the island population, P1a, is particularly vulnerable to climate change and other stochastic factors due to the isolated location and worse genetic health. To prevent the localized extinction of this population, as seen in Finland (Copete & Sharpe, 2020), long-term population and genetic monitoring, as well as habitat protection, should be implemented.

## Acknowledgments

We thank the University of Washington Burke Museum and the University of Minnesota Bell Museum for providing specimen loans. This project was supported by the General Research Fund (University Grants Committee) granted to S.Y.W.S (project no. 17124422). We also thank the computation resources provided by the Information Technology Services at the University of Hong Kong. We also thank Dr. Alison Cloutier for her valuable comments.

## Data Accessibility Statement

The genetic data will be uploaded to the National Center for Biotechnology Information (NCBI). Detailed accessibility information will be provided after the manuscript is accepted.

## Author Contributions

Conceptualization: S.Y.W.S; sample collection: S.Y.W.S. and G.C.; methodology and lab works: G.C. and S.Y.W.S.; analysis: G.C.; visualization: G.C.; writing-original draft: G.C.; writing-review and editing: all authors; supervision, project administration, and funding acquisition: S.Y.W.S.

## Supplemental Information

### Supplementary Figures

**Figure S1.**
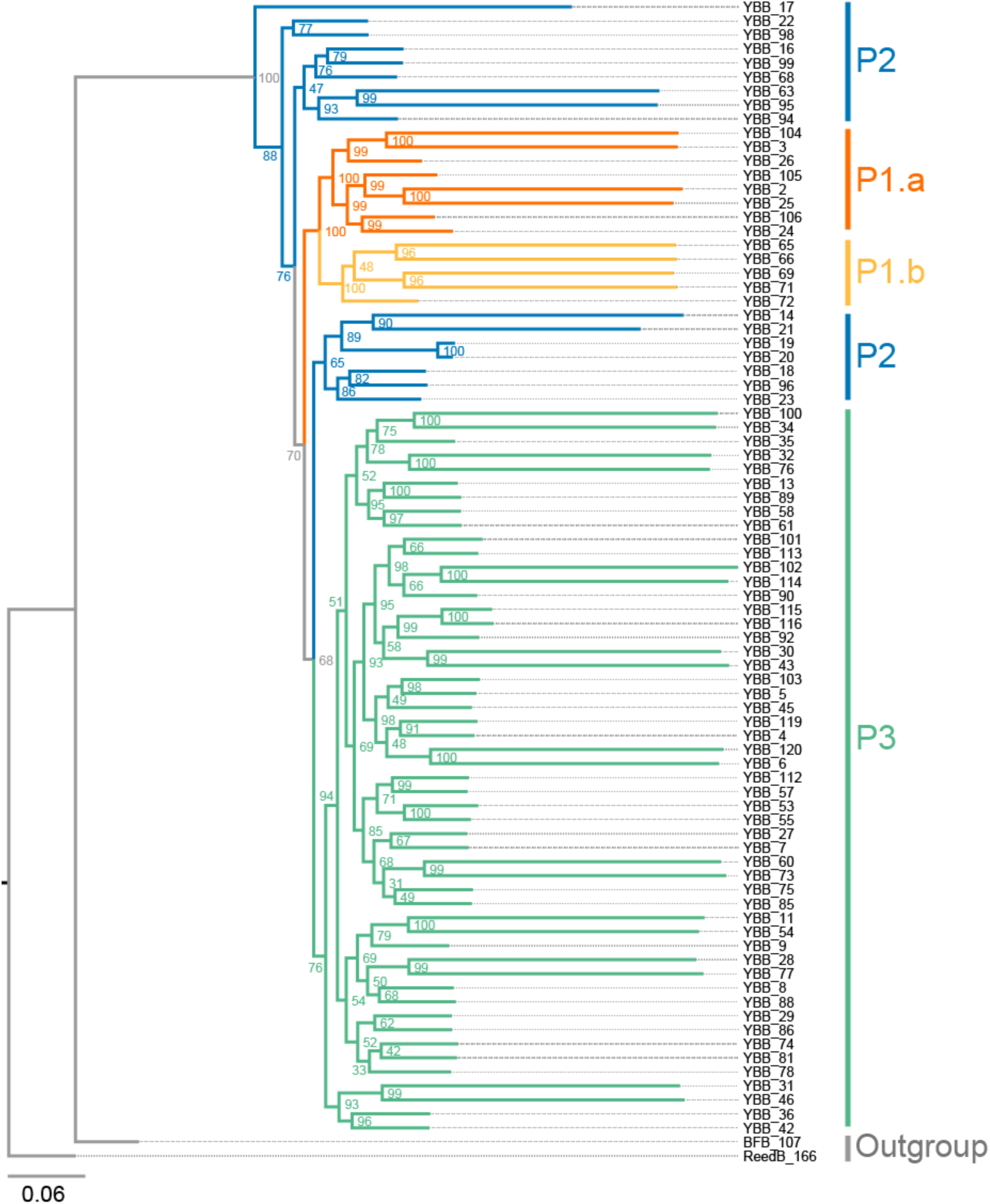
The maximum likelihood tree was inferred using IQ-TREE2 based on the Z-linkedSNPs.

**Figure S2.**
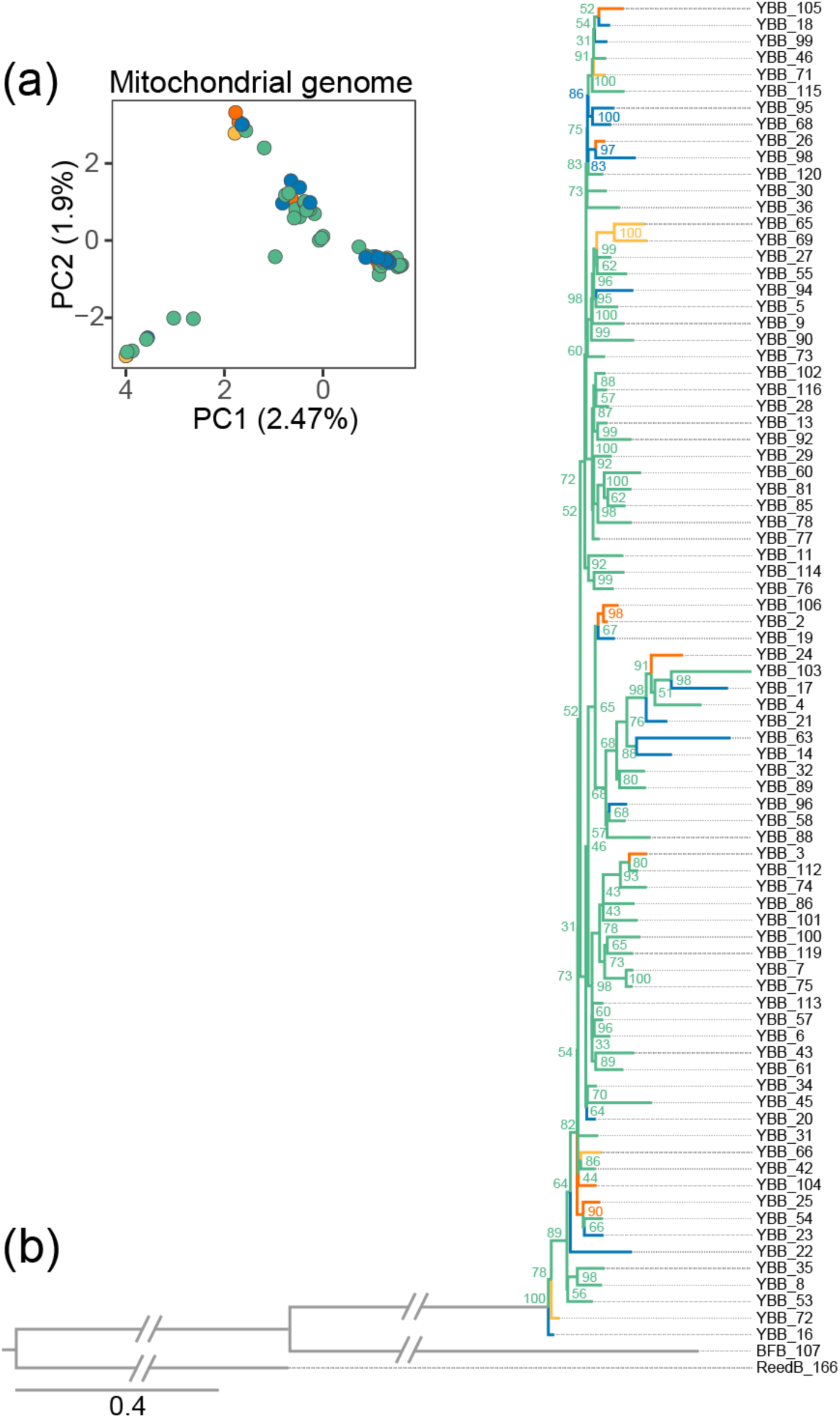
The Principal components analysis (a) and maximum likelihood tree (b) were inferred using mitochondrial data.

**Figure S3.**
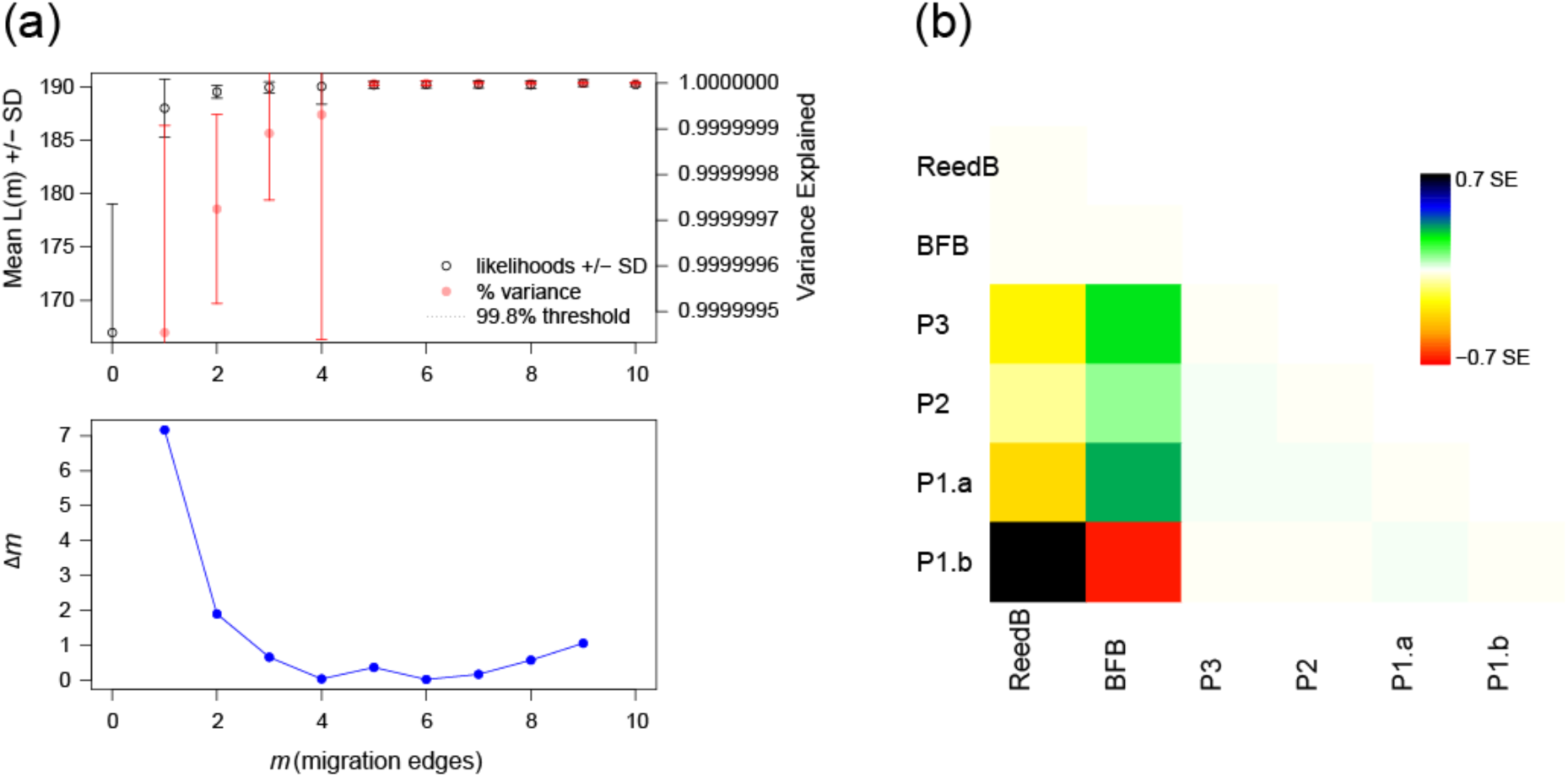
The estimated best migration edges (m) for Treemix. (a) The output of *OptM* with the best migration edges is 1 (m=1). The upper panel indicates the mean and standard deviation (SD) for the composite likelihood (left-y-axis) and the proportion of variance explained (right-y-axis). We have tested migration edges ranging from 0 to 10. The lower panel indicates the second-order rate of change (Δm) based on different migration edges. The highest Δm indicates the best migration edges. (b) The residual fit of each pair of populations based on the maximum likelihood tree inferring using Treemix. The color indicates the number of residuals; residuals larger than zero indicate the candidate for admixture events.

**Figure S4.**
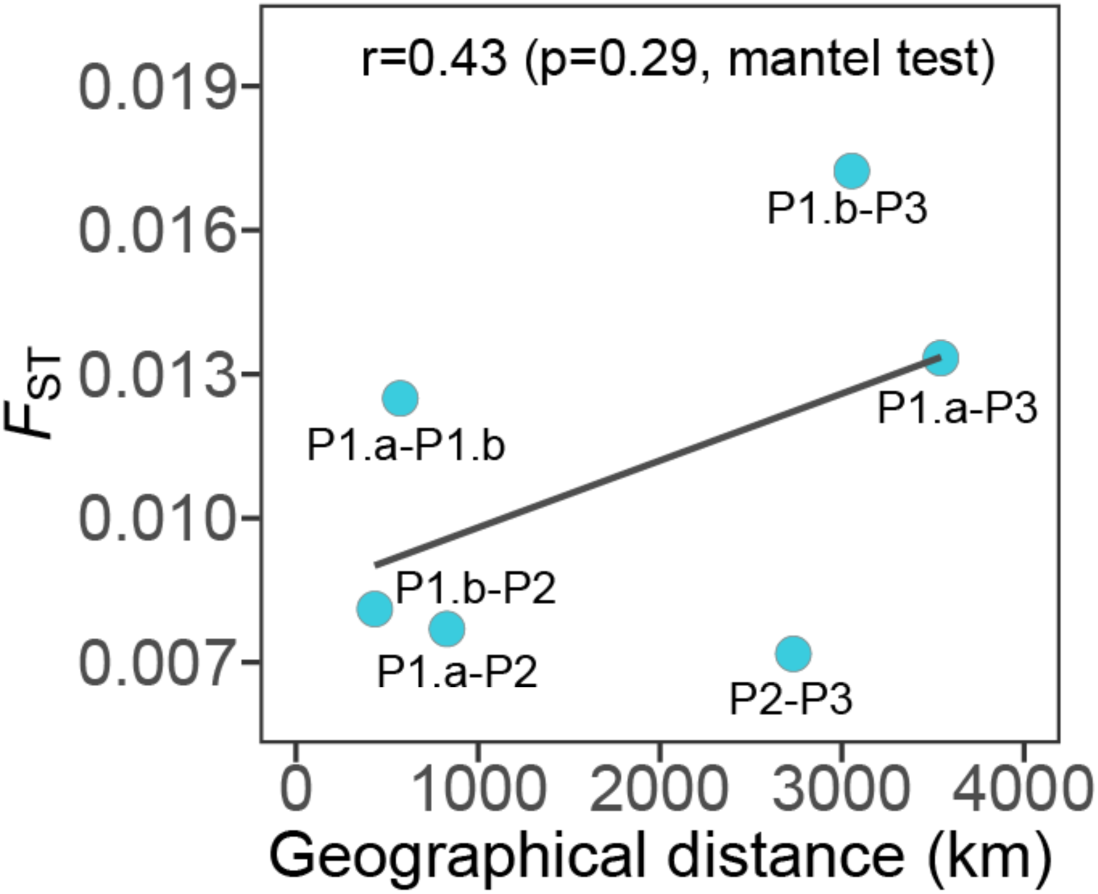
The correlation between the pairwise genetic differentiation (*F*_ST_) and the geographic distance. Each point indicates the *F*_ST_ value between each population pair.

**Figure S5.**
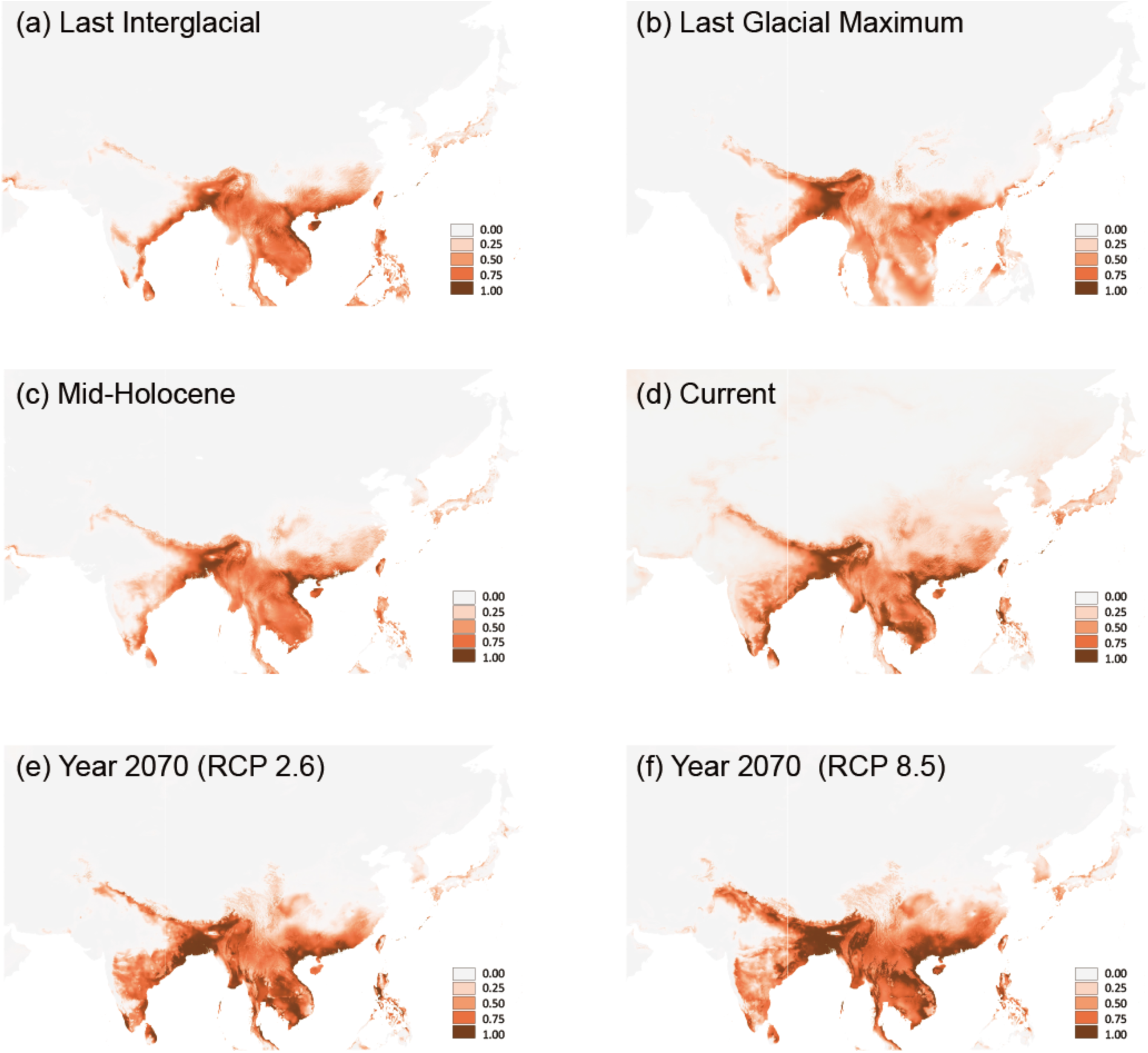
The predicted suitable wintering range of yellow-breasted bunting. Reconstruction of ecological niche models for the wintering area of yellow-breasted buntings in the Last Interglacial (LIG), Last Glacial Maximum (LGM), Mid-Holocene (MH), current, and year 2070 with different Representative Concentration Pathways (RCP2.6 and RCP 8.5). The color indicates the probability of suitable conditions in the models (i.e. darker color indicates more suitable).

**Figure S6.**
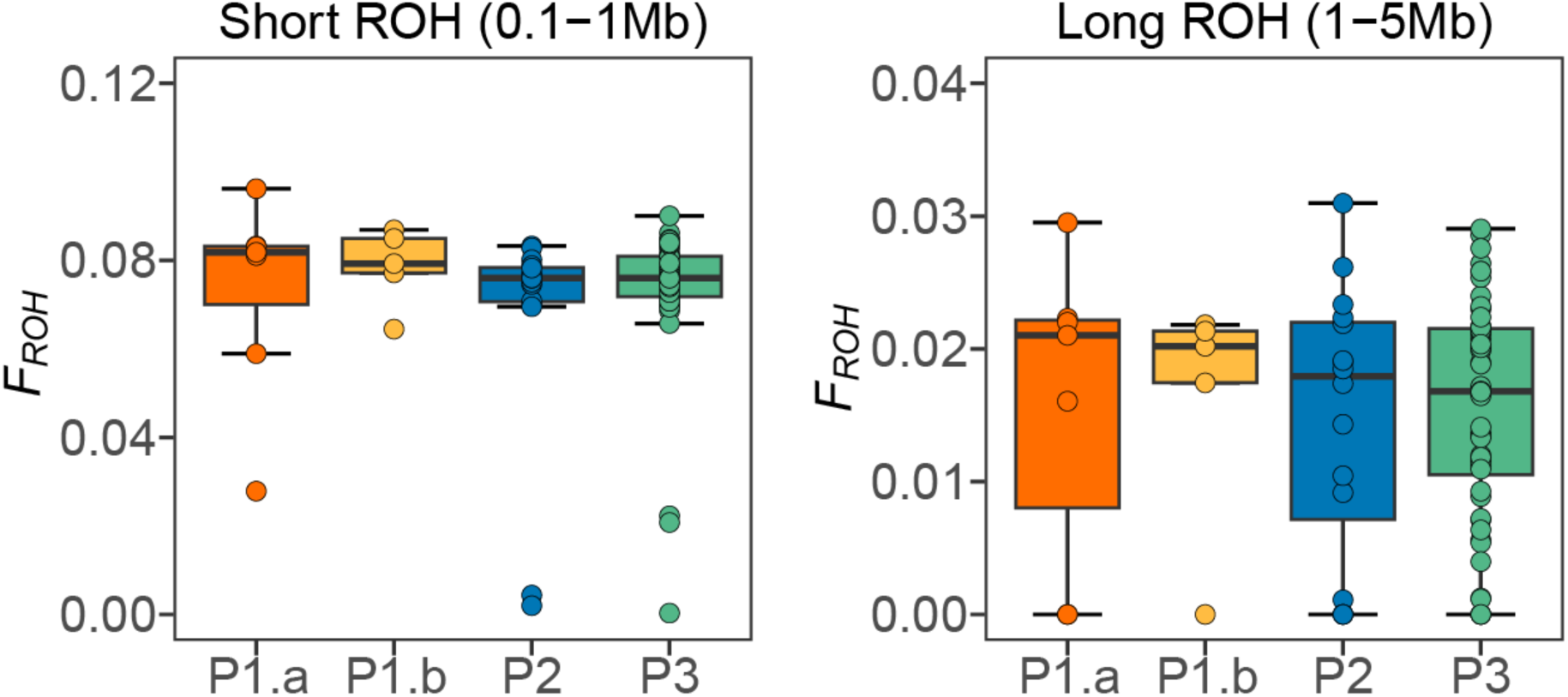
The inbreeding coefficient, *F_ROH_*, was inferred based on different lengths of ROHs. The difference between each pair of populations is not significant.

**Figure S7.**
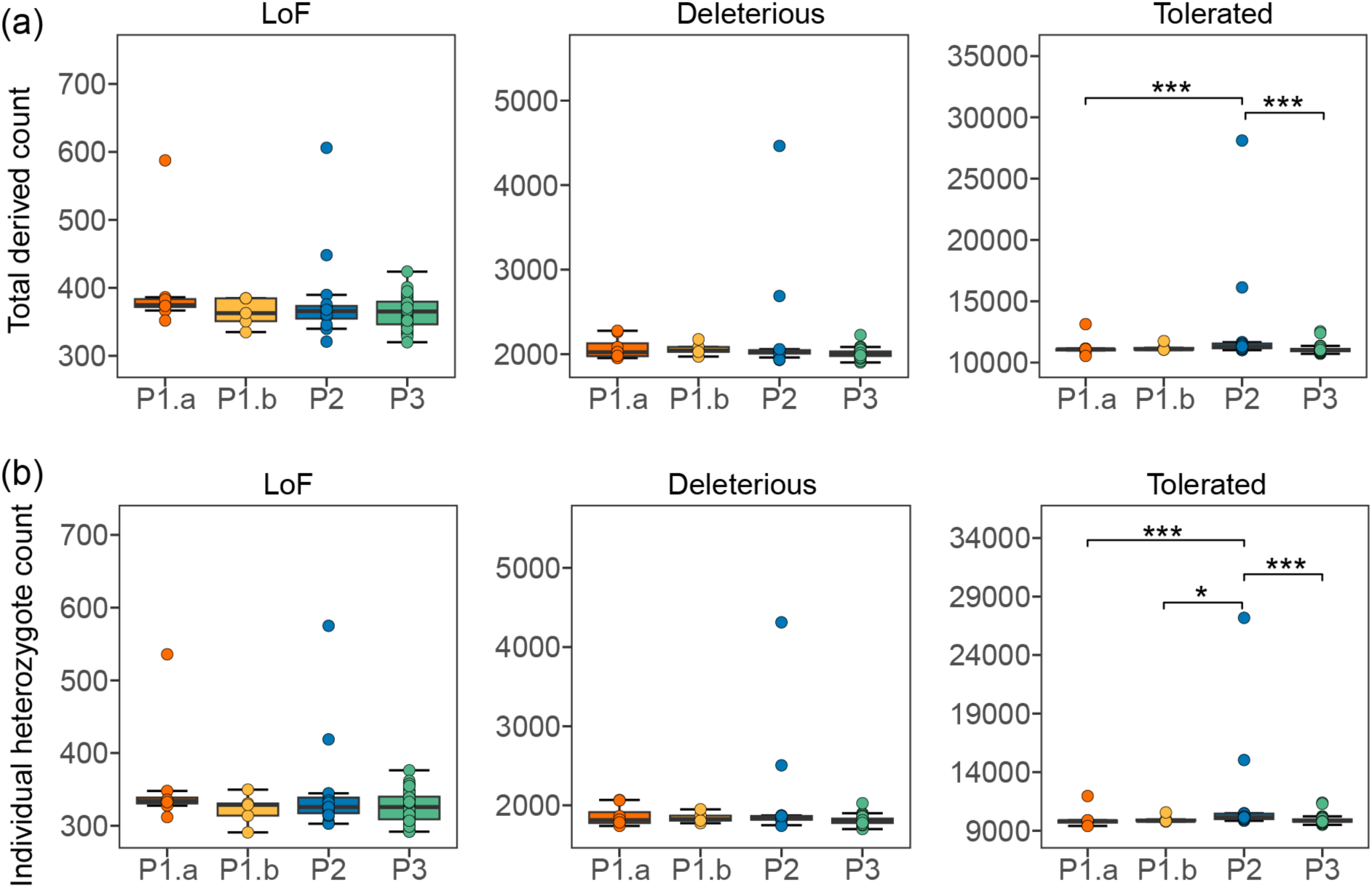
The accumulation of deleterious mutations of yellow-breasted bunting. (a) Total number of LoF, deleterious nonsynonymous, and tolerated nonsynonymous mutations. (b) The number of LoF, deleterious nonsynonymous, and tolerated nonsynonymous heterozygous mutations. The significant differences between population pairs are indicated by asterisks (*). * p<0.05; ** p<0.01; *** p<0.001.

**Table S1.**
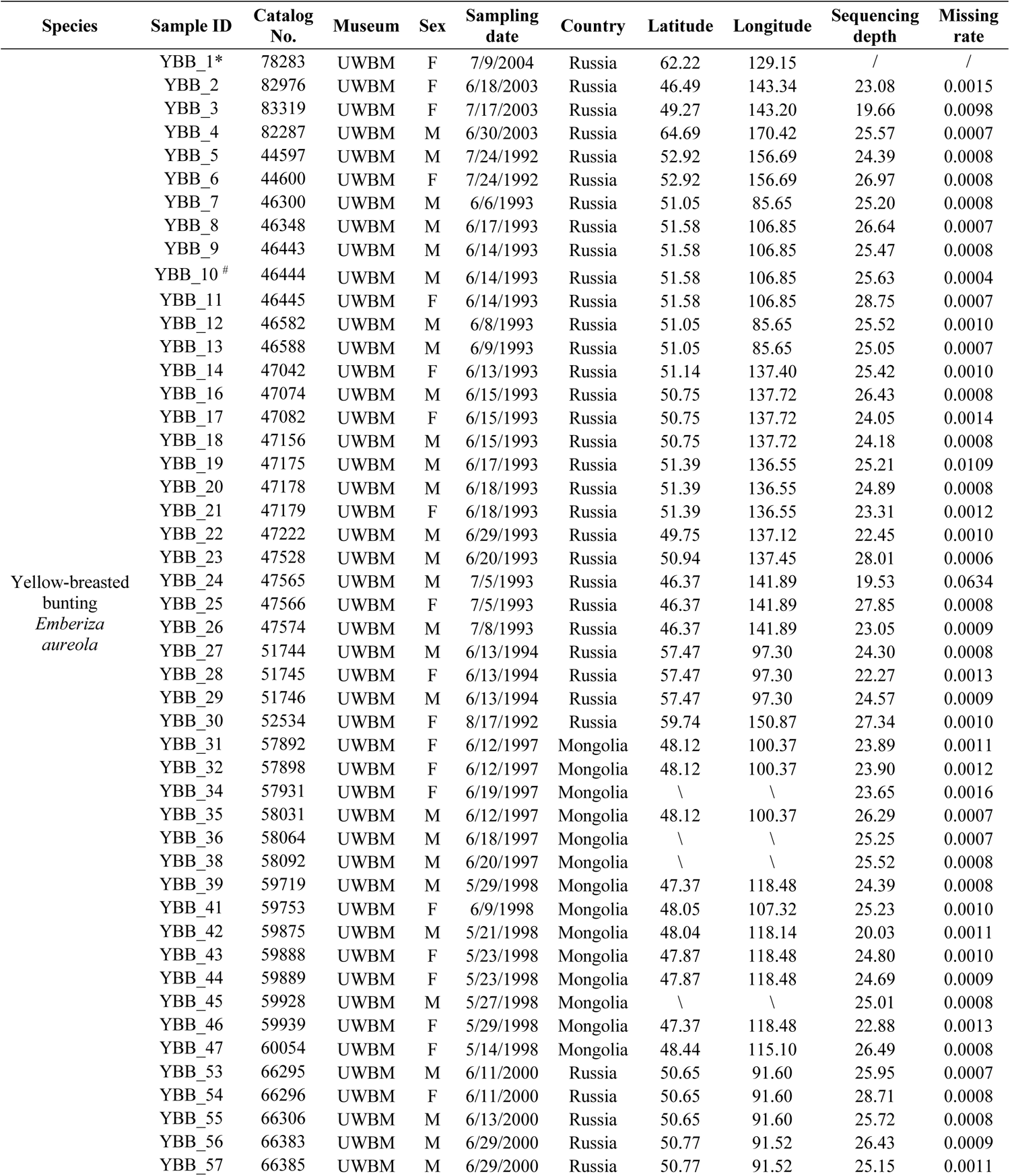
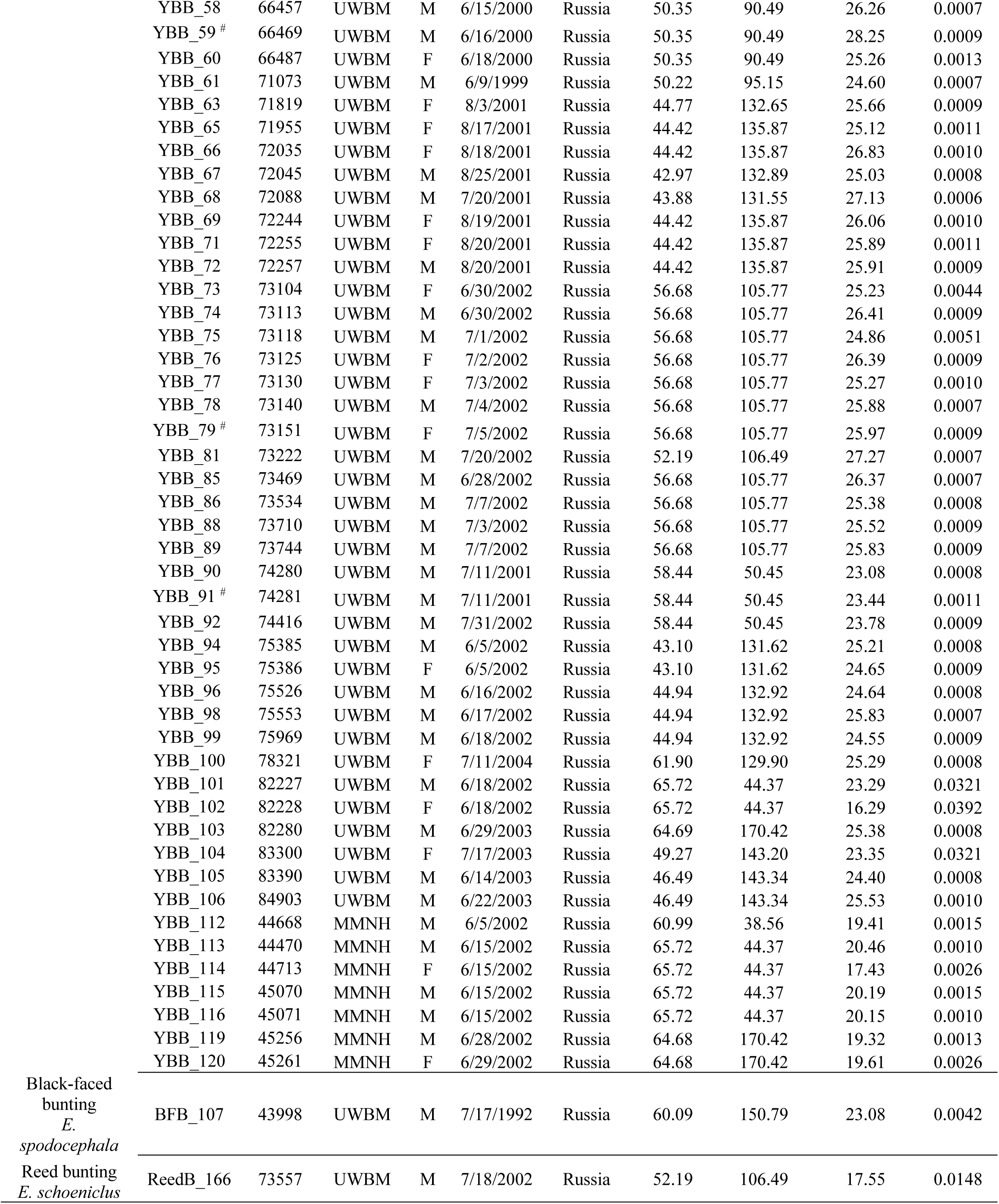
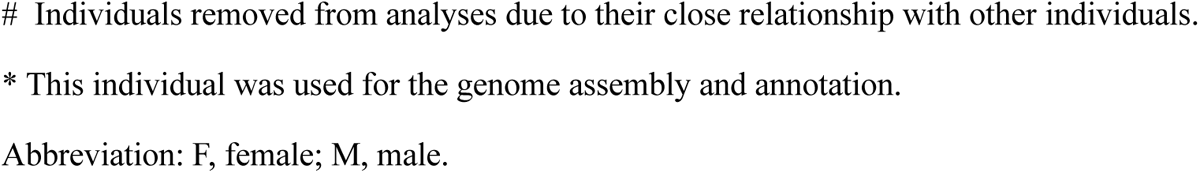
Sampling and sequencing information of three bunting species used in this study.

**Table S2.**
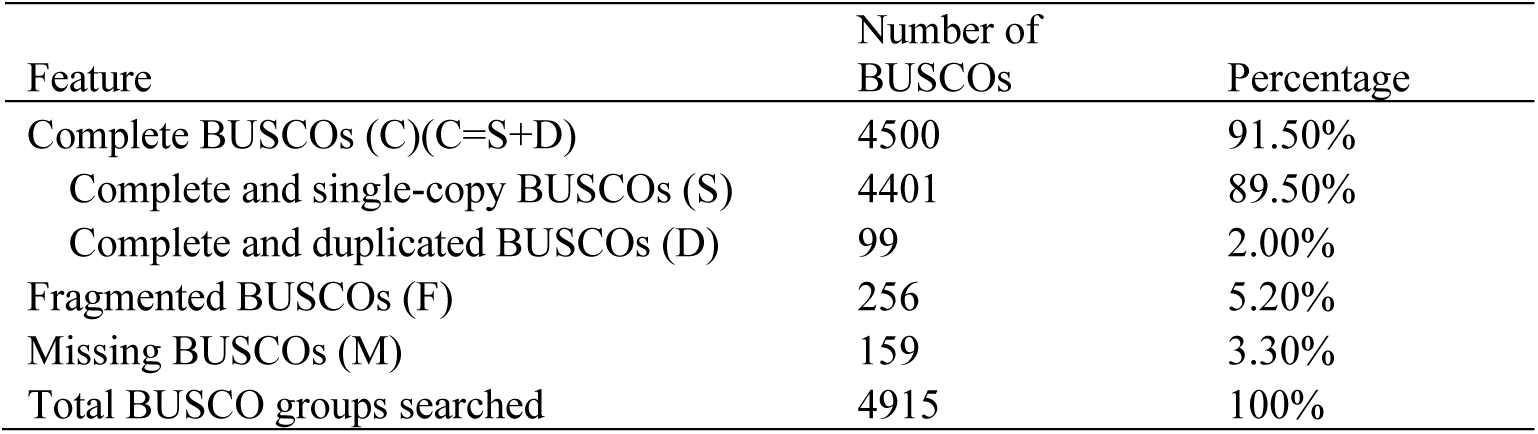
Output from BUSCO analyses to assess the yellow-breasted bunting genome completeness by searching for single-copy orthologs from aves dataset.

**Table S3.**
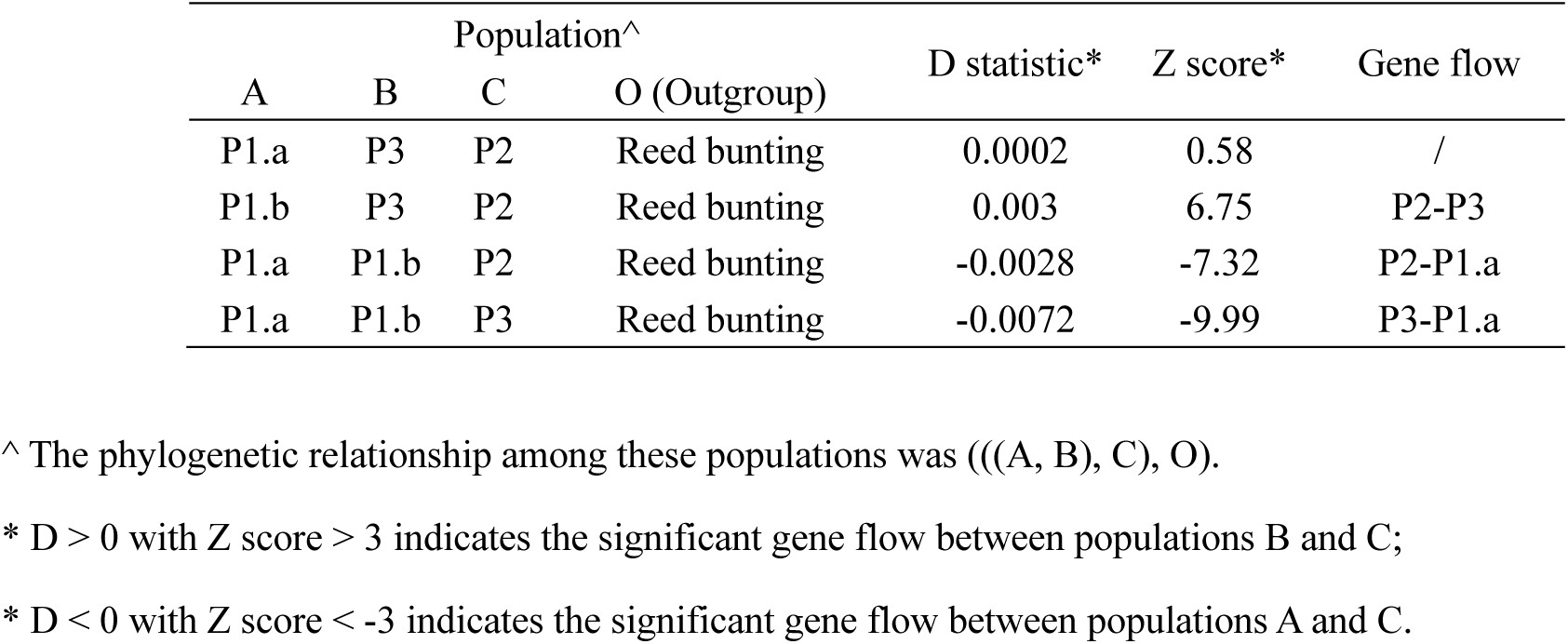
The results of Patterson’s *D* statistic for each trio of yellow-breasted bunting populations.

## References

Albright, T. P., Pidgeon, A. M., Rittenhouse, C. D., Clayton, M. K., Flather, C. H., Culbert, P. D., … Radeloff, V. C. (2010). Effects of drought on avian community structure. Global Change Biology, 16(8), 2158–2170. doi:10.1111/j.1365-2486.2009.02120.x

Alexander, D. H., Novembre, J., & Lange, K. (2009). Fast model-based estimation of ancestry in unrelated individuals. Genome Research, 19(9), 1655–1664. doi:10.1101/gr.094052.109

Andrews, S. (2017). FastQC: a quality control tool for high throughput sequence data. 2010. W29–33.

Backstrom, N., Forstmeier, W., Schielzeth, H., Mellenius, H., Nam, K., Bolund, E., … Ellegren, H. (2010). The recombination landscape of the zebra finch *Taeniopygia guttata* genome. Genome Research, 20(4), 485–495. doi:10.1101/gr.101410.109

Bairoch, A., & Apweiler, R. (2000). The SWISS-PROT protein sequence database and its supplement TrEMBL in 2000. Nucleic Acids Research, 28(1), 45–48. doi:DOI 10.1093/nar/28.1.45

Bertorelle, G., Raffini, F., Bosse, M., Bortoluzzi, C., Iannucci, A., Trucchi, E., … van Oosterhout, C. (2022). Genetic load: genomic estimates and applications in non-model animals. Nature Reviews Genetics, 23(8), 492–503. doi:10.1038/s41576-022-00448-x

BirdLife International. (2024). Species factsheet: Emberiza aureola. Downloaded from https://datazone.birdlife.org/species/factsheet/yellow-breasted-bunting-emberiza-aureola on 07/05/2024.

Blomqvist, D., Pauliny, A., Larsson, M., & Flodin, L. Å. (2010). Trapped in the extinction vortex? Strong genetic effects in a declining vertebrate population. BMC Evol Biol, 10. doi:10.1186/1471-2148-10-33

Ceballos, F. C., Joshi, P. K., Clark, D. W., Ramsay, M., & Wilson, J. F. (2018). Runs of homozygosity: windows into population history and trait architecture. Nature Reviews Genetics, 19(4), 220–234. doi:10.1038/nrg.2017.109

Chen, S. F., Zhou, Y. Q., Chen, Y. R., & Gu, J. (2018). fastp: an ultra-fast all-in-one FASTQ preprocessor. Bioinformatics, 34(17), 884–890. doi:10.1093/bioinformatics/bty560

Chevreux, B., Pfisterer, T., Drescher, B., Driesel, A. J., Müller, W. E. G., Wetter, T., & Suhai, S. (2004). Using the miraEST assembler for reliable and automated mRNA transcript assembly and SNP detection in sequenced ESTs. Genome Research, 14(6), 1147–1159. doi:10.1101/gr.1917404

Cingolani, P., Platts, A., Wang, L. L., Coon, M., Nguyen, T., Wang, L., … Ruden, D. M. (2012). A program for annotating and predicting the effects of single nucleotide polymorphisms, SnpEff: SNPs in the genome of Drosophila melanogaster strain w1118; iso-2; iso-3. *Fly*, 6(2), 80–92. doi:10.4161/fly.19695

Collins, A. F., Bush, M. B., & Sachs, J. P. (2013). Microrefugia and species persistence in the Galapagos highlands: a 26,000-year paleoecological perspective. Front Genet, 4, 269. doi:10.3389/fgene.2013.00269

Copete, J. L., & Sharpe, C. J. (2020). Yellow-breasted Bunting (Emberiza aureola), version 1.0. In Birds of the World (J. del Hoyo, A. Elliott, J. Sargatal, D. A. Christie, and E. de Juana, Editors). Cornell Lab of Ornithology, Ithaca, NY, USA. Retrieved from 10.2173/bow.yebbun.01

Couvet, D. (2002). Deleterious effects of restricted gene flow in fragmented populations. Conservation Biology, 16(2), 369–376. doi:DOI 10.1046/j.1523-1739.2002.99518.x

Cowie, R. H., Bouchet, P., & Fontaine, B. (2022). The Sixth Mass Extinction: fact, fiction or speculation? Biological Reviews, 97(2), 640–663. doi:10.1111/brv.12816

Danecek, P., Auton, A., Abecasis, G., Albers, C. A., Banks, E., DePristo, M. A., … Grp, G. P. A. (2011). The variant call format and VCFtools. Bioinformatics, 27(15), 2156–2158. doi:10.1093/bioinformatics/btr330

De Laet, J., & Summers-Smith, J. D. (2007). The status of the urban house sparrow in north-western Europe. Journal of Ornithology, 148, S275–S278. doi:10.1007/s10336-007-0154-0

Dierickx, E. G., Sin, S. Y. W., van Veelen, H. P. J., Brooke, M. D., Liu, Y., Edwards, S. V., & Martin, S. H. (2020). Genetic diversity, demographic history and neo-sex chromosomes in the Critically Endangered Raso lark. Proceedings of the Royal Society B-Biological Sciences, 287(1922), 20192613. doi:10.1098/rspb.2019.2613

Dixon, P. (2003). VEGAN, a package of R functions for community ecology. Journal of Vegetation Science, 14(6), 927–930. doi:10.1111/j.1654-1103.2003.tb02228.x

Donald, P. F., Green, R. E., & Heath, M. F. (2001). Agricultural intensification and the collapse of Europe’s farmland bird populations. Proceedings of the Royal Society B-Biological Sciences, 268(1462), 25–29. doi:10.1098/rspb.2000.1325

Dussex, N., Van Der Valk, T., Morales, H. E., Wheat, C. W., Díez-del-Molino, D., Von Seth, J., … Phillippy, A. M. (2021). Population genomics of the critically endangered kākāpō. Cell Genomics, 1(1), 100002. doi:10.1016/j.xgen.2021.100002

Fagan, W. F., & Holmes, E. E. (2006). Quantifying the extinction vortex. Ecology Letters, 9(1), 51–60. doi:10.1111/j.1461-0248.2005.00845.x

Fitak, R. R. (2021). OptM: estimating the optimal number of migration edges on population trees using Treemix. Biol Methods Protoc, 6(1), bpab017. doi:10.1093/biomethods/bpab017

Flynn, J. M., Hubley, R., Goubert, C., Rosen, J., Clark, A. G., Feschotte, C., & Smit, A. F. (2020). RepeatModeler2 for automated genomic discovery of transposable element families. Proceedings of the National Academy of Sciences of the United States of America, 117(17), 9451–9457. doi:10.1073/pnas.1921046117

Frankham, R., Ballou, J. D., & Briscoe, D. A. (2010). Introduction to conservation genetics 2nd edn. Cambridge, UK.: Cambridge University Press.

Gandra, M., Assis, J., Martins, M. R., & Abecasis, D. (2021). Reduced Global Genetic Differentiation of Exploited Marine Fish Species. Molecular Biology and Evolution, 38(4), 1402–1412. doi:10.1093/molbev/msaa299

Glemin, S. (2003). How are deleterious mutations purged? Drift versus nonrandom mating. Evolution, 57(12), 2678–2687. doi:10.1111/j.0014-3820.2003.tb01512.x

Grabherr, M. G., Russell, P., Meyer, M., Mauceli, E., Alfoldi, J., Di Palma, F., & Lindblad-Toh, K. (2010). Genome-wide synteny through highly sensitive sequence alignment: Satsuma. Bioinformatics, 26(9), 1145–1151. doi:10.1093/bioinformatics/btq102

Grantham, R. (1974). Amino-acid difference formula to help explain protein evolution. Science, 185(4154), 862–864. doi:10.1126/science.185.4154.862

Grossen, C., Guillaume, F., Keller, L. F., & Croll, D. (2020). Purging of highly deleterious mutations through severe bottlenecks in Alpine ibex. Nature Communications, 11(1), 1001. doi:10.1038/s41467-020-14803-1

Hahn, C., Bachmann, L., & Chevreux, B. (2013). Reconstructing mitochondrial genomes directly from genomic next-generation sequencing reads-a baiting and iterative mapping approach. Nucleic Acids Research, 41(13), e129. doi:10.1093/nar/gkt371

Haller, B. C., & Messer, P. W. (2023). SLiM 4: Multispecies Eco-Evolutionary Modeling. American Naturalist, 201(5), 127–139. doi:10.1086/723601

Hannon, G. J. (2010). Fastx-toolkit. Downloaded from http://hannonlab.cshl.edu/fastx_toolkit/.

Hedrick, P. W., & Garcia-Dorado, A. (2016). Understanding inbreeding depression, purging, and genetic rescue. Trends in Ecology & Evolution, 31(12), 940–952. doi:10.1016/j.tree.2016.09.005

Heim, W., Chan, S., Hölzel, N., Ktitorov, P., Mischenko, A., & Kamp, J. (2021). East Asian buntings: Ongoing illegal trade and encouraging conservation responses. Conservation Science and Practice, 3(6), e405. doi:10.1111/csp2.405

Hewitt, G. M. (2004). Genetic consequences of climatic oscillations in the Quaternary. Philosophical Transactions of the Royal Society B-Biological Sciences, 359(1442), 183–195. doi:10.1098/rstb.2003.1388

Hijmans, R. J., Williams, E., Vennes, C., & Hijmans, M. R. J. (2017). Package ‘geosphere’. Spherical trigonometry, 1(7), 1–45.

Holm, S. R., & Svenning, J. C. (2014). 180,000 Years of Climate Change in Europe: Avifaunal Responses and Vegetation Implications. PLoS One, 9(4), e94021. doi:10.1371/journal.pone.0094021

Holt, C., & Yandell, M. (2011). MAKER2: an annotation pipeline and genome-database management tool for second-generation genome projects. Bmc Bioinformatics, 12, 1–14. doi:10.1186/1471-2105-12-491

Huerta-Cepas, J., Szklarczyk, D., Heller, D., Hernández-Plaza, A., Forslund, S. K., Cook, H., … Bork, P. (2019). eggNOG 5.0: a hierarchical, functionally and phylogenetically annotated orthology resource based on 5090 organisms and 2502 viruses. Nucleic Acids Research, 47(D1), D309–D314. doi:10.1093/nar/gky1085

Hung, C. M., Shaner, P. J. L., Zink, R. M., Liu, W. C., Chu, T. C., Huang, W. S., & Li, S. H. (2014). Drastic population fluctuations explain the rapid extinction of the passenger pigeon. Proceedings of the National Academy of Sciences of the United States of America, 111(29), 10636–10641. doi:10.1073/pnas.1401526111

Huynh, S., Cloutier, A., Chen, G. L., Chan, D. T. C., Lam, D. K., Huyvaert, K. P., … Sin, S. Y. W. (2023). Whole-genome Analyses Reveal Past Population Fluctuations and Low Genetic Diversities of the North Pacific Albatrosses. Molecular Biology and Evolution, 40(7), msad155. doi:10.1093/molbev/msad155

Inger, R., Gregory, R., Duffy, J. P., Stott, I., Vorisek, P., & Gaston, K. J. (2015). Common European birds are declining rapidly while less abundant species’ numbers are rising. Ecology Letters, 18(1), 28–36. doi:10.1111/ele.12387

Jombart, T. (2008). adegenet: a R package for the multivariate analysis of genetic markers. Bioinformatics, 24(11), 1403–1405. doi:10.1093/bioinformatics/btn129

Jombart, T., & Ahmed, I. (2011). adegenet 1.3-1: new tools for the analysis of genome-wide SNP data. Bioinformatics, 27(21), 3070–3071. doi:10.1093/bioinformatics/btr521

Kalinowski, S. T. (2011). The computer program STRUCTURE does not reliably identify the main genetic clusters within species: simulations and implications for human population structure. Heredity, 106(4), 625–632. doi:10.1038/hdy.2010.95

Kamp, J., Oppel, S., Ananin, A. A., Durnev, Y. A., Gashev, S. N., Hölzel, N., … Chan, S. (2015). Global population collapse in a superabundant migratory bird and illegal trapping in China. Conservation Biology, 29(6), 1684–1694. doi:10.1111/cobi.12537

Kardos, M., Armstrong, E. E., Fitzpatrick, S. W., Hauser, S., Hedrick, P. W., Miller, J. M., … Funk, W. C. (2021). The crucial role of genome-wide genetic variation in conservation. Proceedings of the National Academy of Sciences of the United States of America, 118(48), e2104642118. doi:10.1073/pnas.2104642118

Kardos, M., Taylor, H. R., Ellegren, H., Luikart, G., & Allendorf, F. W. (2016). Genomics advances the study of inbreeding depression in the wild. Evolutionary Applications, 9(10), 1205–1218. doi:10.1111/eva.12414

Kardos, M., Zhang, Y. L., Parsons, K. M., Yunga, A., Kang, H., Xu, X., … Li, S. H. (2023). Inbreeding depression explains killer whale population dynamics. Nature Ecology & Evolution, 7(5), 675–686. doi:10.1038/s41559-023-01995-0

Keilwagen, J., Hartung, F., & Grau, J. (2019). GeMoMa: Homology-Based Gene Prediction Utilizing Intron Position Conservation and RNA-seq Data. Methods Mol Biol, 1962, 161–177. doi:10.1007/978-1-4939-9173-0_9

Khan, A., Patel, K., Shukla, H., Viswanathan, A., van der Valk, T., Borthakur, U., … Ramakrishnan, U. (2021). Genomic evidence for inbreeding depression and purging of deleterious genetic variation in Indian tigers. Proceedings of the National Academy of Sciences of the United States of America, 118(49), e2023018118. doi:10.1073/pnas.2023018118

Kyriazis, C. C., Beichman, A. C., Brzeski, K. E., Hoy, S. R., Peterson, R. O., Vucetich, J. A., … Wayne, R. K. (2023). Genomic Underpinnings of Population Persistence in Isle Royale Moose. Molecular Biology and Evolution, 40(2), msad021. doi:10.1093/molbev/msad021

Kyriazis, C. C., Wayne, R. K., & Lohmueller, K. E. (2021). Strongly deleterious mutations are a primary determinant of extinction risk due to inbreeding depression. Evolution Letters, 5(1), 33–47. doi:10.1002/evl3.209

Lam, D. K., Frantz, A. C., Burke, T., Geffen, E., & Sin, S. Y. W. (2023). Both selection and drift drive the spatial pattern of adaptive genetic variation in a wild mammal. Evolution, 77(1), 221–238. doi:10.1093/evolut/qpac014

Li, H., & Durbin, R. (2009). Fast and accurate short read alignment with Burrows-Wheeler transform. Bioinformatics, 25(14), 1754–1760. doi:10.1093/bioinformatics/btp324

Li, H., & Durbin, R. (2011). Inference of human population history from individual whole-genome sequences. Nature, 475(7357), 493–496. doi:10.1038/nature10231

Li, H., Handsaker, B., Wysoker, A., Fennell, T., Ruan, J., Homer, N., … Proc, G. P. D. (2009). The Sequence Alignment/Map format and SAMtools. Bioinformatics, 25(16), 2078–2079. doi:10.1093/bioinformatics/btp352

Li, S. B., Li, B., Cheng, C., Xiong, Z. J., Liu, Q. B., Lai, J. H., … Yan, J. Q. (2014). Genomic signatures of near-extinction and rebirth of the crested ibis and other endangered bird species. Genome Biology, 15(12), 1–17. doi:10.1186/S13059-014-0557-1

Liu, X. M., & Fu, Y. X. (2015). Exploring population size changes using SNP frequency spectra. Nature Genetics, 47(5), 555–559. doi:10.1038/ng.3254

Liu, Y. C., Schröder, J., & Schmidt, B. (2013). Musket: a multistage k-mer spectrum based error corrector for Illumina sequence data. Bioinformatics, 29(3), 308–315. doi:10.1093/bioinformatics/bts690

Malinsky, M., Matschiner, M., & Svardal, H. (2021). Dsuite - Fast-statistics and related admixture evidence from VCF files. Molecular Ecology Resources, 21(2), 584–595. doi:10.1111/1755-0998.13265

Manichaikul, A., Mychaleckyj, J. C., Rich, S. S., Daly, K., Sale, M., & Chen, W. M. (2010). Robust relationship inference in genome-wide association studies. Bioinformatics, 26(22), 2867–2873. doi:10.1093/bioinformatics/btq559

McKenna, A., Hanna, M., Banks, E., Sivachenko, A., Cibulskis, K., Kernytsky, A., … DePristo, M. A. (2010). The Genome Analysis Toolkit: A MapReduce framework for analyzing next-generation DNA sequencing data. Genome Research, 20(9), 1297–1303. doi:10.1101/gr.107524.110

Mingay, G. E. (1977). The agricultural revolution: changes in agriculture, 1650–1880.

Minh, B. Q., Schmidt, H. A., Chernomor, O., Schrempf, D., Woodhams, M. D., von Haeseler, A., & Lanfear, R. (2020). IQ-TREE 2: New Models and Efficient Methods for Phylogenetic Inference in the Genomic Era. Molecular Biology and Evolution, 37(5), 1530–1534. doi:10.1093/molbev/msaa015

Murray, G. G. R., Soares, A. E. R., Novak, B. J., Schaefer, N. K., Cahill, J. A., Baker, A. J., … Shapiro, B. (2017). Natural selection shaped the rise and fall of passenger pigeon genomic diversity. Science, 358(6365), 951–954. doi:10.1126/science.aao0960

Nadachowska-Brzyska, K., Li, C., Smeds, L., Zhang, G. J., & Ellegren, H. (2015). Temporal dynamics of avian populations during Pleistocene revealed by whole-genome sequences. Current Biology, 25(10), 1375–1380. doi:10.1016/j.cub.2015.03.047

Pan, T., Ren, L., Zhu, X., Yan, L., Hu, C., Chang, Q., & Zhang, B. (2015). Mitochondrial genome of the *Emberiza aureola* (Emberizidae: *Emberiza*). Mitochondrial DNA, 26(1), 121–122. doi:10.3109/19401736.2013.814112

Park, J.-G., Park, C.-U., Jin, K.-S., Kim, Y.-M., Kim, H.-Y., Jeong, S.-Y., & Nam, D.-H. (2020). Moult and plumage patterns of the critically endangered Yellow-breasted Bunting (Emberiza aureola) at a stopover site in Korea. Journal of ornithology, 161(1), 257–266. doi:10.1007/s10336-019-01716-0

Patterson, N., Moorjani, P., Luo, Y. T., Mallick, S., Rohland, N., Zhan, Y. P., … Reich, D. (2012). Ancient Admixture in Human History. Genetics, 192(3), 1065–1093. doi:10.1534/genetics.112.145037

Phillips, S. J., Anderson, R. P., & Schapire, R. E. (2006). Maximum entropy modeling of species geographic distributions. Ecological Modelling, 190(3-4), 231–259. doi:10.1016/j.ecolmodel.2005.03.026

Picard. (2019). Picard Toolkit. GitHub Repository. http://broadinstitute.github.io/picard/; Broad Institute.

Pickrell, J. K., & Pritchard, J. K. (2012). Inference of Population Splits and Mixtures from Genome-Wide Allele Frequency Data. Plos Genetics, 8(11), 1–1. doi:10.1371/journal.pgen.1002967

Price, M. N., Dehal, P. S., & Arkin, A. P. (2010). FastTree 2-Approximately Maximum-Likelihood Trees for Large Alignments. PLoS One, 5(3), e9490. doi:10.1371/journal.pone.0009490

Purcell, S., Neale, B., Todd-Brown, K., Thomas, L., Ferreira, M. A. R., Bender, D., … Sham, P. C. (2007). PLINK: A tool set for whole-genome association and population-based linkage analyses. American Journal of Human Genetics, 81(3), 559–575. doi:10.1086/519795

Rigal, S., Dakos, V., Alonso, H., Aunins, A., Benko, Z., Brotons, L., … Devictor, V. (2023). Farmland practices are driving bird population decline across Europe. Proceedings of the National Academy of Sciences of the United States of America, 120(21), e2216573120. doi:10.1073/pnas.2216573120

Robinson, J. A., Kyriazis, C. C., Nigenda-Morales, S. F., Beichman, A. C., Rojas-Bracho, L., Robertson, K. M., … Morin, P. A. (2022). The critically endangered vaquita is not doomed to extinction by inbreeding depression. Science, 376(6593), 635–639. doi:10.1126/science.abm1742

Robinson, J. A., Raikkonen, J., Vucetich, L. M., Vucetich, J. A., Peterson, R. O., Lohmueller, K. E., & Wayne, R. K. (2019). Genomic signatures of extensive inbreeding in Isle Royale wolves, a population on the threshold of extinction. Science Advances, 5(5), eaau0757. doi:10.1126/sciadv.aau0757

Santiago, E., Novo, I., Pardinas, A. F., Saura, M., Wang, J. L., & Caballero, A. (2020). Recent demographic history inferred by high-resolution analysis of linkage disequilibrium. Molecular Biology and Evolution, 37(12), 3642–3653. doi:10.1093/molbev/msaa169

Schorger, A. W. (1995). The Passenger Pigeon: Its Natural History and Extinction: University of Wisconsin Press, Madison, WI.

Simao, F. A., Waterhouse, R. M., Ioannidis, P., Kriventseva, E. V., & Zdobnov, E. M. (2015). BUSCO: assessing genome assembly and annotation completeness with single-copy orthologs. Bioinformatics, 31(19), 3210–3212. doi:10.1093/bioinformatics/btv351

Simonsen, M., Mailund, T., & Pedersen, C. N. (2008). Rapid neighbour-joining. In Algorithms in Bioinformatics: 8th International Workshop, WABI 2008, Karlsruhe, Germany, September 15-19, 2008. Proceedings 8 (pp. 113-122). Springer Berlin Heidelberg.

Sin, S. Y. W., Hoover, B. A., Nevitt, G. A., & Edwards, S. V. (2021). Demographic History, Not Mating System, Explains Signatures of Inbreeding and Inbreeding Depression in a Large Outbred Population. American Naturalist, 197(6), 658–676. doi:10.1086/714079

Smeds, L., Qvarnström, A., & Ellegren, H. (2016). Direct estimate of the rate of germline mutation in a bird. Genome Research, 26(9), 1211–1218. doi:10.1101/gr.204669.116

Smit, A. F. A., Hubley, R., & Green, P. (2020). “RepeatMasker” downloaded from http://www.repeatmasker.org.

Smith, H. G., Ryegård, A., & Svensson, S. (2012). Is the large-scale decline of the starling related to local changes in demography? Ecography, 35(8), 741–748. doi:10.1111/j.1600-0587.2011.06310.x

Snyder, C. W. (2016). Evolution of global temperature over the past two million years. Nature, 538(7624), 226–228. doi:10.1038/nature19798

Tamada, K., Hayama, S., Umeki, M., Takada, M., & Tomizawa, M. (2017). Drastic declines in Brown Shrike and Yellow-breasted Bunting at the Lake Utonai Bird Sanctuary, Hokkaido. Ornithological Science, 16(1), 51–57. doi:10.2326/osj.16.51

Tamada, K., Tomizawa, M., Umeki, M., & Takada, M. (2014). Population trends of grassland birds in Hokkaido, focussing on the drastic decline of the Yellow-breasted Bunting. Ornithological Science, 13(1), 29–40. doi:10.2326/osj.13.29

Tataru, P., & Bataillon, T. (2019). polyDFEv2.0: testing for invariance of the distribution of fitness effects within and across species. Bioinformatics, 35(16), 2868–2869. doi:10.1093/bioinformatics/bty1060

Terhorst, J., Kamm, J. A., & Song, Y. S. (2017). Robust and scalable inference of population history from hundreds of unphased whole genomes. Nature Genetics, 49(2), 303–309. doi:10.1038/ng.3748

Thompson, F. M. L. (1968). The second agricultural revolution, 1815-1880. The Economic History Review 21(1), 62–77. doi:10.2307/2592204

Tsukada, M. (1983). Vegetation and Climate during the Last Glacial Maximum in Japan. Quaternary Research, 19(2), 212–235. doi:10.1016/0033-5894(83)90006-6

Välimäki, K., Lindén, A., & Lehikoinen, A. (2016). Velocity of density shifts in Finnish landbird species depends on their migration ecology and body mass. Oecologia, 181(1), 313–321. doi:10.1007/s00442-015-3525-x

van Vuuren, D. P., Edmonds, J., Kainuma, M., Riahi, K., Thomson, A., Hibbard, K., … Rose, S. K. (2011). The representative concentration pathways: an overview. Climatic Change, 109(1), 5–31. doi:10.1007/s10584-011-0148-z

Wang, I. J. (2013). Examining the Full Effects of Landscape Heterogeneity on Spatial Genetic Variation: A Multiple Matrix Regression Approach for Quantifying Geographic and Ecological Isolation. Evolution, 67(12), 3403–3411. doi:10.1111/evo.12134

Wang, I. J., Glor, R. E., & Losos, J. B. (2013). Quantifying the roles of ecology and geography in spatial genetic divergence. Ecology Letters, 16(2), 175–182. doi:10.1111/ele.12025

Wang, P. C., Hou, R., Wu, Y., Zhang, Z. W., Que, P. J., & Chen, P. (2022). Genomic status of yellow-breasted bunting following recent rapid population decline. Iscience, 25(7). doi:10.1016/j.isci.2022.104501

Xu, H. B., Luo, X., Qian, J., Pang, X. H., Song, J. Y., Qian, G. R., … Chen, S. L. (2012). FastUniq: A fast de novo duplicates removal tool for paired short reads. PLoS One, 7(12), e52249. doi:10.1371/journal.pone.0052249

Zhan, X. J., Pan, S. K., Wang, J. Y., Dixon, A., He, J., Muller, M. G., … Bruford, M. W. (2013). Peregrine and saker falcon genome sequences provide insights into evolution of a predatory lifestyle. Nature Genetics, 45(5), 563–U142. doi:10.1038/ng.2588

Zheng, G. X. Y., Lau, B. T., Schnall-Levin, M., Jarosz, M., Bell, J. M., Hindson, C. M., … Ji, H. P. (2016). Haplotyping germline and cancer genomes with high-throughput linked-read sequencing. Nature Biotechnology, 34(3), 303–311. doi:10.1038/nbt.3432

Zheng, Y. C., & Janke, A. (2018). Gene flow analysis method, the D-statistic, is robust in a wide parameter space. Bmc Bioinformatics, 19, 10. doi:10.1186/s12859-017-2002-4

